# Single-cell analysis of an engineered organoid-based model of pancreatic cancer identifies hypoxia as a contributing factor in the determination of transcriptional subtypes

**DOI:** 10.1101/2024.03.05.583412

**Authors:** Natalie Landon-Brace, Brendan T. Innes, Simon Latour, Jose L. Cadavid, Ileana L. Co, Cassidy M. Tan, Ferris Nowlan, Sybil Drissler, Faiyaz Notta, Hartland Warren Jackson, Gary D. Bader, Alison P. McGuigan

## Abstract

Pancreatic ductal adenocarcinoma (PDAC) is a high-mortality cancer characterized by its aggressive, treatment-resistant phenotype and a complex tumour microenvironment (TME) featuring significant hypoxia. Bulk transcriptomic analysis has identified the “classical” and “basal-like” transcriptional subtypes which have prognostic value in PDAC; however, it remains unclear how microenvironmental heterogeneity contributes to the expression of these transcriptional signatures. Here, we used single cell transcriptome analysis of the organoid TRACER platform to explore the effect of oxygen and other microenvironmental gradients on PDAC organoid cells. We found that the microenvironmental gradients present in TRACER significantly impact the distribution of organoid transcriptional phenotypes and the enrichment of gene sets linked to cancer progression and treatment resistance. More significantly, we found that microenvironmental gradients drive changes in the expression of the classical and basal-like transcriptional subtype gene signatures. This effect is likely dominated by the oxygen gradients in TRACER, as hypoxia alone induced decreases in the expression of classical marker GATA6 at both the gene and protein level in PDAC cells. This work suggests that hypoxia contributes to determining transcriptional subtypes in PDAC and broadly underscores the importance of considering microenvironmental gradients in organoid-based transcriptomic studies of PDAC.

## Introduction

Pancreatic ductal adenocarcinoma (PDAC), the most common type of pancreatic cancer, is a high-mortality cancer with a five-year survival rate of approximately 11%^1^. As most patients present with metastatic disease and are therefore ineligible for curative surgery, the mainstay of treatment is chemotherapy, which offers only a modest survival benefit^2^. There is significant heterogeneity in responses to treatment between patients, with most experiencing some response to chemotherapy before the disease ultimately progresses^3^. Genomic studies of PDAC indicate that the majority of tumours exhibit recurrent mutations in KRAS, CDKN2A, TP53 and SMAD4, suggesting that clinical heterogeneity likely arises from non-genetic sources^3,4^. Accordingly, understanding the non-genetic factors that contribute to PDAC progression and response to therapy is essential for patient stratification of disease sub-type and the identification of more effective treatment strategies.

Transcriptomic analysis of PDAC has revealed several subtype classification schemes intended to clarify clinical heterogeneity^4–8^. The most robust classification scheme based on tumour cell-specific gene expression broadly subdivides PDAC into “basal-like” and “classical” tumours^7,8^. The basal-like subtype can be distinguished by its low GATA6 expression and is associated with more aggressive disease with reduced responsiveness to chemotherapy and poorer patient survival^7,9–12^. Both classical and basal-like tumour cells can co-exist in the same tumour, with some evidence suggesting that the more aggressive basal-like cells arise as a subpopulation of cells in an otherwise classical tumour^3,13^. This heterogeneity has been associated with more advanced stages of PDAC and poorer survival compared to homogenously classical tumours^14^. Though some genetic factors that contribute to the determination of transcriptional subtypes have been identified^3,13^, the influence of the tumour microenvironment (TME), a critical player in PDAC, remains unclear.

The TME in PDAC is characterized by a dense desmoplastic stroma, an immunosuppressive milieu, and significant hypoxia that serves both pro- and anti-tumorigenic function and contributes to therapeutic resistance^15–18^. Hypoxia, a common feature of many solid tumours, is particularly pronounced in PDAC as the characteristic desmoplastic stroma can impair vascularization and perfusion, which exacerbates the low oxygen conditions arising from rapid tumour growth ^18–20^. Notably, hypoxia can contribute to tumour cell invasion and metastasis, metabolic reprogramming, survival, and chemoresistance^10,18,19^. Furthermore, hypoxia can activate fibroblasts in the TME enhancing the fibrotic reaction and creating a positive feedback loop that further perpetuates the hypoxic environment^18,19^. While hypoxia has been associated with basal-like disease^10,21^, this relationship was identified using bulk sequencing of patient tumour samples. Thus, it is unclear whether hypoxia causes tumour cells to adopt a more basal-like phenotype and how this relationship may be influenced by other cell types present in the TME. Additionally, bulk analysis masks heterogeneity in cellular responses to microenvironmental conditions. Accordingly, further investigation is needed to characterize the relationship between hypoxia and the transcriptional subtypes in PDAC and to explore the effect of tumour-microenvironment interactions on tumour cell heterogeneity in this context.

Recent advances in experimental techniques and *in vitro* modelling tools offer a unique avenue for exploring this interaction. Single cell RNA-sequencing (scRNA-seq) has enabled high-resolution characterization of the heterogeneity of the PDAC TME and tumour cell phenotypes^3,14,22^. However, scRNA-seq is challenging in PDAC due to low tumour cellularity and the limited availability of tumour samples from patients with more advanced disease^3,23^. Additionally, scRNA-seq inherently requires removal of tumour cells from the TME, limiting the ability to correlate transcriptomic features with specific microenvironmental features. While spatial transcriptomic retain the spatial relationships between cells in the TME, commercial tools have not yet reached single-cell resolution and the presence of other cell types present in the TME can confound tumour cell intrinsic responses to microenvironmental conditions^24,25^.

To overcome these limitations, ex vivo models can be used to study tumour heterogeneity in defined microenvironments. Patient-derived organoids (PDOs) generated from PDAC tumours can be used to expand tumour cells ex vivo to increase cellularity for sequencing and improve access to patient material from across the disease spectrum while retaining key genomic and phenotypic features of patient tumours^14,26–29^. However, PDOs, if grown as small clusters of cells, lack microenvironmental heterogeneity and in larger organoids cells cannot be easily retrieved from specific locations in the organoid structure. These issues limit the extent to which organoid cultures can be used to probe the impact of TME features on cell phenotypes^30,31^. To enable modelling of more aspects of the TME using organoids, our group previously described the “Tissue Roll for Analysis of Cellular Environment and Response” (TRACER), a tumour organoid model that can be disassembled to study cellular responses to a 3D microenvironment featuring gradients of oxygen and other small molecules^32–34^. In TRACER, PDOs are dissociated into single organoid cells, embedded in a hydrogel and infiltrated into a cellulose paper strip that is rolled around an oxygen-impermeable core. This creates a six-layer stacked construct that features cell consumption-derived microenvironmental gradients and recapitulates several phenotypic changes associated with tumour hypoxia^32,33^. Importantly, TRACERs can be disassembled and segmented into individual layers for analysis, allowing spatial correlation of organoid cell phenotypes with their location in the microenvironmental gradient^32,33^. Thus, TRACER offers a unique platform for investigating specific interactions between tumour intrinsic heterogeneity and microenvironmental factors such as oxygen gradients.

Here, we exploited the ability for controlled disassembly of TRACER and performed scRNA-seq of organoid cell populations isolated from individual TRACER layers to investigate the impact of microenvironmental gradients on transcriptional heterogeneity. We identified 7 organoid cell transcriptional phenotypes that were present in varying proportions across the six TRACER layers. Interestingly, these phenotypes were not equally sensitive to the microenvironmental gradients present in TRACER. Our data also show that classical and basal-like organoid cells co-exist in TRACER and that the proportion of basal-like organoid cells increases in the inner layers of TRACER, likely driven predominantly by hypoxia. We also validated our RNAseq results in TRACERs fabricated with PDOs derived from additional patient samples. Thus, our study broadly underscores the impact of microenvironmental gradients on heterogeneity in the tumour cell population in PDAC.

## Results

### scRNA-seq of TRACER reveals 7 organoid cell transcriptional phenotypes

To explore the transcriptional heterogeneity of tumour cells in a microenvironmental gradient, we fabricated TRACER constructs with organoid cells from a Stage IIB PDAC PDO model (PPTO.46). Organoid cells were cultured in TRACER for 24 h before being sectioned into individual layers for scRNA-seq. This timepoint was selected to allow for comparison with our previous work on organoid cell phenotypes from the same PDO source^32^. A schematic of the experimental design is shown in **Fig. 1A**. Transcriptional information was obtained from a total of 10,978 cells from across all 6 layers after quality control, with a median of 2,018 retained cells per layer (range: 1,362-2,154 cells per layer) and a median of 4,420 genes detected per cell (**SI Fig. 1A-D**). All organoid cells were confirmed to be malignant ductal cells by expression of characteristic genes (**SI Fig. 1E**)^14^. To investigate the transcriptional features of the organoid cells across all layers of TRACER, we performed k-nearest neighbour (kNN) clustering and identified 7 clusters after assessing for robustness (**Fig. 1B, Fig. S2A-E**). We also observed that layer of origin was not the primary determinant of cell clustering (**Fig. 1C**). This suggested that transcriptional heterogeneity exists both within and between the layers of TRACER.

**Figure 1:**
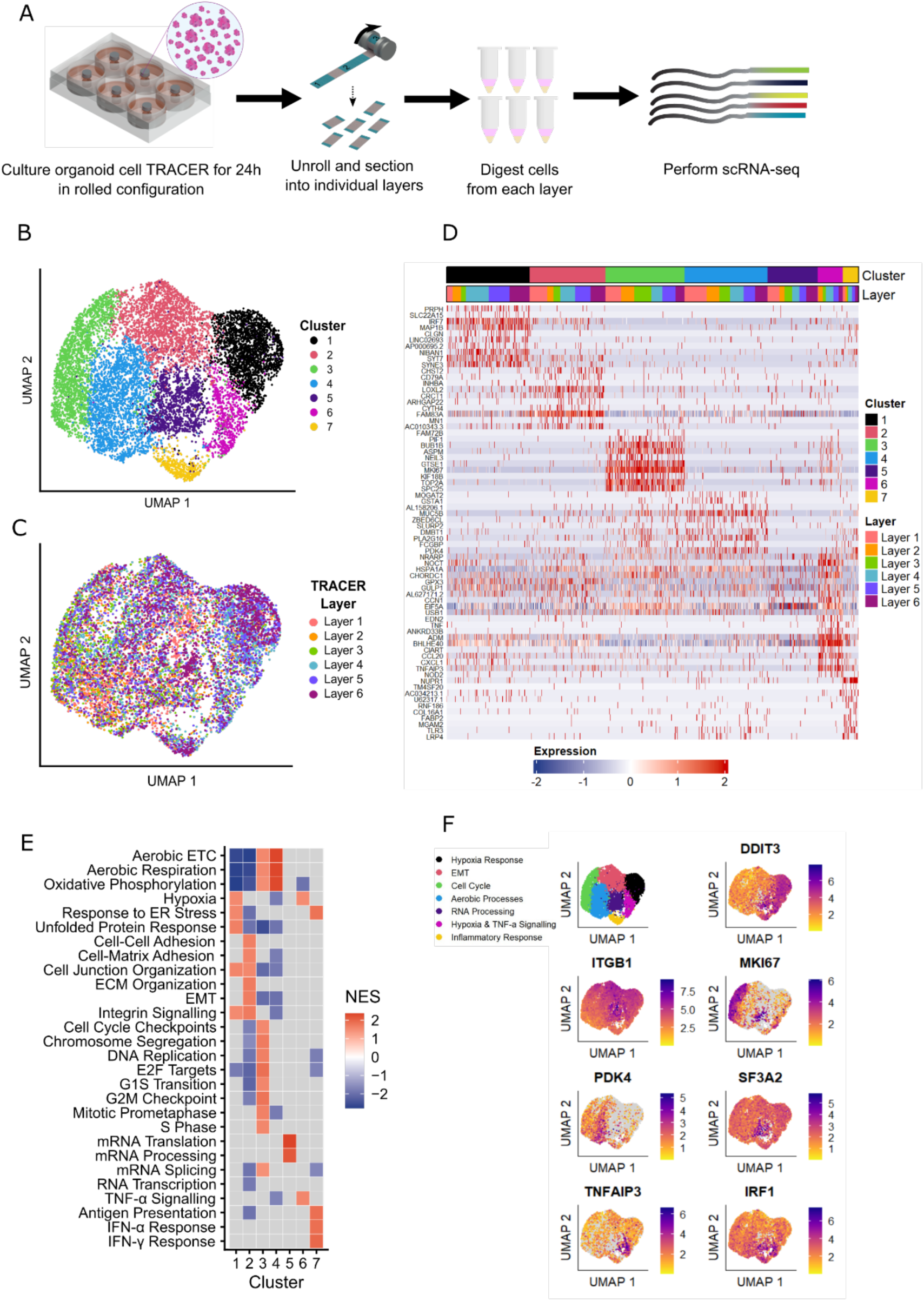
Transcriptomic analysis of organoid cells isolated from TRACER reveals heterogeneity that is not exclusively dependent on layer of origin. (A) Schematic of the experimental workflow. (B) UMAP plot of organoid cells isolated from all layers of the TRACER construct. Seven organoid cell clusters were identified. (C) UMAP plot of organoid cells isolated from all layers of TRACER coloured by layer of origin reveals cluster identity was determined by gene expression patterns rather than exclusively layer of origin. (D) Heatmap of 10 most differentially expressed genes for each of the seven single-cell clusters. Cluster identity and layer of origin are shown in the coloured bars. (E) Gene set enrichment analysis by cluster using gene sets compiled from multiple databases indicating gene sets that were significantly enriched (adjusted p-value < 0.05) across databases. The maximum normalized enrichment score (NES) for the reoccurring gene sets is represented. Gray boxes indicate no significant enrichment. A summary of the gene sets and their source database is provided in Table S1. (F) Visualization of a representative gene for each cluster-defining gene set supports cluster annotation [DDIT3 – Unfolded Protein Response, Hypoxia (Hypoxia Response); ITGB1 – EMT, Integrin Signalling (EMT); MKI67 – Cell proliferation, E2F targets (Cell Cycle); PDK4 – Oxidative Phosphorylation (Aerobic Processes); SF3A2 – mRNA Processing (RNA Processing); TNFAIP3 – TNFα signalling (Hypoxia & TNF-α Signalling); IRF1 – Interferon signalling (Inflammatory Response)].

We next explored the cellular phenotypes represented in these clusters by identifying cluster-specific marker genes (**Fig. 1D**) and performing gene set enrichment analysis (GSEA) using gene sets compiled from several databases^17^. Clusters were identified based on the enrichment of common gene sets across databases (**Fig. 1E, SI Table 1**). We identified two clusters as representing the tumor cell response to hypoxia (clusters 1 – “hypoxia response” – and 6), with cluster 6 additionally showing enriched expression of genes related to TNF-α signalling (“hypoxia & TNF-α signalling”). Cluster 2 represented cells with enriched expression of genes related to the epithelial-mesenchymal transition (EMT), cell-cell and cell-matrix adhesion, ECM organization, and integrin signalling (“EMT”). Cluster 3 (“cell cycle”) represented cells progressing through the cell cycle with enrichment of cell cycle checkpoint and transition genes. Cluster 4 represented cells with enriched expression of genes related to aerobic respiration, oxidative phosphorylation, and the electron transport chain (“aerobic processes”). Cluster 5 represented cells with increased expression of genes related to mRNA processing and translation (“RNA processing”). However, few genes were significantly positively enriched in this cluster, suggesting that these cells were predominantly defined by the absence of other specific marker genes. Finally, cluster 7 represented cells with increased expression of genes related to antigen presentation and Type I interferon (IFN) response and signalling (“inflammatory response”). Cluster annotations were verified by visualizing the expression of a representative gene from the cluster-defining gene set(s) (**Fig. 1F**). These organoid cell subpopulations broadly align with enriched pathways reported in previous sequencing studies of PDAC and PDAC organoid cells^3,6,14,22,26,35,36^. Importantly, our data reflect the expected transcriptional response to hypoxia, which is a critical factor influencing PDAC tumour cell phenotypes *in vivo*^10,18^ but is largely overlooked in the study of organoids *in vitro*. Our results are also consistent with our previous work demonstrating that TRACER recapitulates relevant aspects of disease biology^32,33^ and emphasize that several subpopulations of organoid cells co-exist across the layers of TRACER.

### Expected and novel gradients in gene expression are observed across the layers of TRACER

Having identified 7 organoid cell clusters in TRACER, we next examined the representation of these clusters in individual layers to explore how organoid cell gene expression patterns varied with the microenvironmental gradients in TRACER. We found that all the identified clusters were represented in each of the TRACER layers, although the proportion of cells in each cluster varied across layers (**Figs. 2A-B**). The proportion of organoid cells in the hypoxia response and the hypoxia response/TNF-α signalling clusters increased from Layer 1 (outer) to Layer 6 (inner), consistent with the hypoxic gradient previously shown to be present in TRACER (linear trend in proportions, p < 0.0001; Table S2)^32^. Conversely, the proportion of organoid cells in the aerobic processes and cell cycle clusters decreased from Layer 1 (outer) to Layer 6 (inner) (linear trend in proportions, p < 0.0001), in line with expectations^32,33^. Surprisingly, the proportion of organoid cells in the inflammatory response cluster increased slightly from Layer 1 (outer) to Layer 6 (inner) (linear trend in proportions, p < 0.01). Previous studies of other cancers have suggested that hypoxia typically downregulates genes associated with the Type I interferon response that were enriched in this cluster^37,38^. However, recent studies have shown that a subset of PDAC tumours exhibit enrichment of IFN signalling associated with metabolic alterations; therefore, IFN signalling is not entirely understood in this context^35,39^. The relationship between hypoxia and the expression of antigen presentation-related genes enriched in this cluster remains poorly understood^40^. The proportion of cells in the EMT cluster significantly differed between the layers (test for equality of proportions, p < 0.0001), although no significant graded trend was observed across the layers (linear trend in proportions, p > 0.05). There was also no significant trend in the proportion of organoid cells in the RNA processing cluster (linear trend in proportions, p > 0.05), suggesting cells characterized by these clusters may exhibit transcriptional phenotypes that are less sensitive to the variations in microenvironmental conditions present in TRACER. This potentially aligns with previous work showing common gene expression patterns between primary and metastatic lesions, which suggests that the activation of some gene programs is conserved in tumour cells that experience different microenvironmental conditions^10,14,41^.

**Figure 2:**
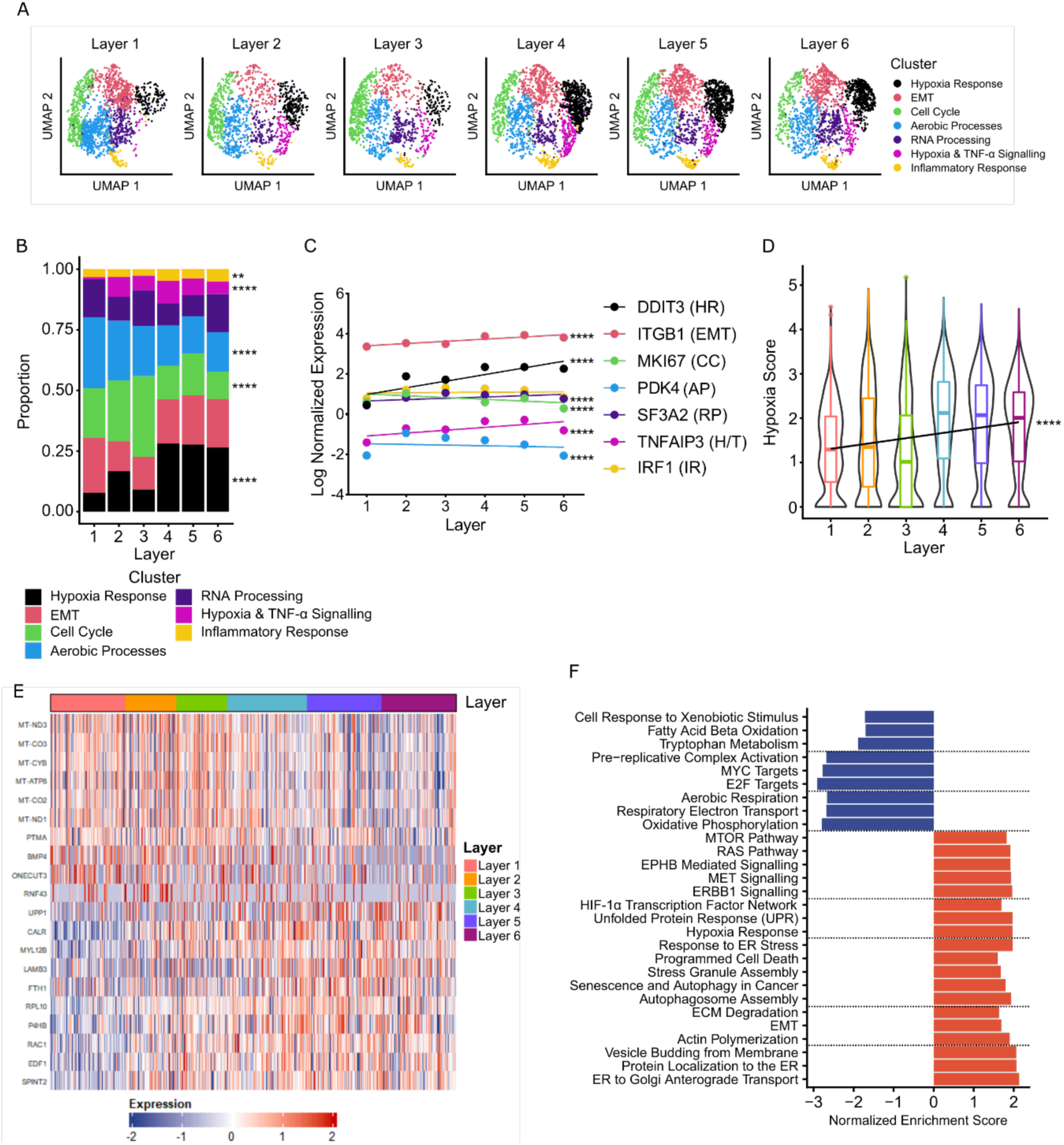
Microenvironmental gradients affect the distribution of organoid cell clusters in TRACER and reveal gene sets with graded expression patterns. (A) UMAP plots showing the distribution of the seven organoid cell subpopulation clusters identified in Figure 1 for each of the layers of TRACER. (B) Quantification of the proportion of cells from each cluster present in each layer. Variations in the cluster proportions across the layers are consistent with expected responses to the hypoxic gradients that are known to exist across the TRACER layers (Chi-square Test for Trend in Proportions, **p < 0.01, ****p < 0.0001). (C) Log normalized expression of cluster-representative genes (from Fig. 1F) show gradients in expression for most cluster markers across the TRACER layers, excluding IRF1 (inflammatory response) (mean ± SEM; slope of the best line vs. 0, ****p < 0.0001). (D) Connor et al. PDAC hypoxia score^3,45^ calculated for the entire organoid cell population from each layer of TRACER. The hypoxia score increases across the layers consistent with the expected graded transcriptional response to the oxygen gradient in TRACER (slope of the best fit line vs. 0, ****p < 0.0001). (E) Heatmap of the top 10 genes that correlate most positively or negatively with layer number. Layer 1 (outer) to Layer 6 (inner) in TRACER shows robust trends with heterogeneity in individual layers. (F) Normalized enrichment score (NES) for selected gene sets significantly enriched (p < 0.05) for correlation with TRACER layer number, where a positive NES (red) indicates increasing enrichment from Layer 1 (outer) to Layer 6 (inner) and a negative NES (blue) indicates decreasing enrichment from Layer 1 to Layer 6. GSEA revealed expected enrichment of gene sets related to cell cycle progression, aerobic metabolism and hypoxia response. Gene sets related to autophagy, protein localization and transport, and several major signalling pathways were also enriched.

Next, to further characterize the relationship between the organoid cell clusters and the microenvironmental gradient in TRACER, we examined trends in the expression of cluster-representative genes (**Fig. 1F**) across the layers of TRACER. Our analysis found that the expression of all representative marker genes excluding IRF1 (inflammatory response) were significantly graded from Layer 1 (outer) to Layer 6 (inner) (slope of the best fit line, p < 0.0001) (**Fig. 2C, Table S2**). Graded trends in DDIT3 (hypoxia response), MKI67 (cell cycle), PDK4 (aerobic processes) and TNFAIP3 (hypoxia response & TNF-α) were similar to the graded trends in cluster proportion and consistent with the expected responses to the oxygen gradient present across the TRACER layers^32^. Interestingly, the graded expression of ITGB1 (EMT), SF3A2 (RNA processing) and IRF1 (inflammatory response) differed from the graded trend observed in the corresponding cluster proportions. This suggests that while individual marker genes are sufficient to capture the most robust aspects of the transcriptional response to the oxygen gradients in TRACER, comprehensive analysis with scRNA-seq is valuable for understanding the more nuanced aspects of microenvironment-dependent reprogramming in the rolled culture as broader transcriptional reprogramming appears to occur in the microenvironmental gradient present in TRACER. Taken together, the distribution of the organoid cell clusters and gene expression patterns we observed across the TRACER layers were consistent with the presence of expected microenvironment-sensitive transcriptional responses, such as increasing hypoxia response, but also suggested the presence of additional transcriptional patterns that were relatively microenvironment-insensitive across the TRACER layers.

Having confirmed that the cluster proportions and cluster marker gene transcription patterns observed across the TRACER layers were consistent with the presence of an oxygen gradient specifically, we next evaluated the cellular response to hypoxia more comprehensively using a hypoxia score inferred from sequencing of PDAC tumours. To do this, we calculated the hypoxia score for each cell as defined by Connor et al. based on the difference in median expression of hypoxia response genes used to stratify PDAC transcriptomes as this was found to correlate with PDAC prognosis and response to therapy^21^. We observed a graded increase in the hypoxia score from Layer 1 (outer) to Layer 6 (inner) (**Fig. 2D**, Table S2; slope of the best fit line vs. 0, p < 0.0001), consistent with the expected response to the oxygen gradient present in TRACER^32^. Interestingly, given that the presence of a hypoxic gradient has previously been empirically confirmed in TRACER, the robust relationship between the hypoxia score and layer number suggests that the Connor et al. hypoxia score likely captures the hypoxic status of tumour cells in vivo effectively.

Confident that our data set reflected previously reported global transcriptional responses to hypoxia, we next identified all gene sets with graded expression patterns across the TRACER layers, independent of cell clustering. To identify gene sets with the greatest enrichment across the layers, we first performed GSEA on the list of genes ranked by the slope of the best fit line for the average expression across TRACER layers (**Fig. S3**). Gene sets related to hypoxia, UPR, endoplasmic reticulum (ER) stress, and autophagy were significantly positively enriched, whereas gene sets related to cell cycle, electron transport, and oxidative phosphorylation were significantly negatively enriched. This was consistent with the gene expression trends previously observed in TRACER and with expected cellular responses to hypoxia, reinforcing the findings of our clustering analysis and suggesting that the cellular response to increasing hypoxia is the most robust change across the TRACER layers^20,32,33,42–44^. Interestingly, several gene sets related to protein localization and ER-Golgi transport were also positively enriched. Though there is some evidence of a Golgi stress response and the upregulation of key Golgi-associated genes in cancer is implicated in metastasis and invasion, it remains unclear in the literature how Golgi function is impacted by microenvironmental conditions^45–51^. Such investigations have been hindered in part due to the lack of appropriate experimental models^45^; therefore, this may be an interesting area for future investigation in TRACER, though beyond the scope of the current work.

In addition to assessing the strongest linear gradients in TRACER, we also explored gene sets with non-linear trends in expression across the layers. To do this we correlated gene expression with TRACER layer number and identified the most positively or negatively correlated genes, suggestive of a consistent trend in gene expression in response to the microenvironmental gradient from Layer 1 (outer) to Layer 6 (inner) (top 10 genes shown in **Fig. 2E**). The expression of these most positively and negatively correlated genes varied within each layer, consistent with the varying proportions of the organoid cell clusters in each of the TRACER layers. GSEA performed on the list of genes ranked by their correlation coefficient (**Fig. 2F, Table S3**) revealed significant negative enrichment of gene sets related to cell cycle progression, aerobic respiration, tryptophan metabolism and fatty acid oxidation, suggesting layer-dependent downregulation of these gene sets. In addition, gene sets related to EMT, hypoxia and ER stress response, and several major signalling pathways showed significant positive enrichment, suggesting layer-dependent upregulation of these gene sets. Building on the expected changes in hypoxia and stress response gene expression, these results align with previous work regarding hypoxia-induced activation of EMT and changes in cell metabolism^20,33,52–54^. Furthermore, previous work in TRACER fabricated with an ovarian cancer cell line showed layer-dependent changes in the concentration of tryptophan and fatty acid metabolites, consistent with our GSEA results; this finding therefore suggests that microenvironmental gradients of metabolites and other molecules are likely captured in organoid TRACER in addition to the robust oxygen gradient^43^. Taken together, broad transcriptional changes occur in organoid cells across the TRACER layers, likely promoting adaptations in a variety of key cellular functions. Furthermore, our data support the notion that this response is activated as a consequence of microenvironmental conditions, in contrast to previous work suggesting that hypoxia response gene activation is an inherent feature of PDAC tumour cells^43^.

### Microenvironmental gradients in TRACER are associated with changes in the expression of transcriptional subtype marker gene sets

Having established that many important transcriptional programs exhibited graded variation across the TRACER layers, we next examined the variation in transcriptional subtypes across the layers. To do this, we first scored the organoid cells in each layer for their expression of the Moffit subtype gene signature. A greater proportion of organoid cells from the hypoxic inner layers of TRACER scored as basal-like compared to those in the outer layers, with an approximately linear increase in the proportion of basal-like cells across the layers (**Figs. 3A-B, S4A**)^3,52^. This increase in the proportion of basal-like organoid cells in the inner layers of TRACER was observed when we repeated our analysis using other basal-like/classical gene signatures identified in recent work (**Figs. S5A-D, S6A-D**)^3,6^. Given the association between the basal-like gene signature and response to gemcitabine therapy, we next examined the predicted sensitivity to gemcitabine across the TRACER layers using a PDO-derived score^3,55^. We found that the predicted gemcitabine sensitivity of the organoid cells decreased from Layer 1 (outer) to Layer 6 (inner) (slope of the best fit line vs. 0, p < 0.0001; **Fig 3C, Table S2**), consistent with our previous work showing organoid cells in the inner layers of TRACER exhibit a decreased response to gemcitabine^32^. This further supports the enrichment of a basal-like transcriptional phenotype across the layers.

**Figure 3:**
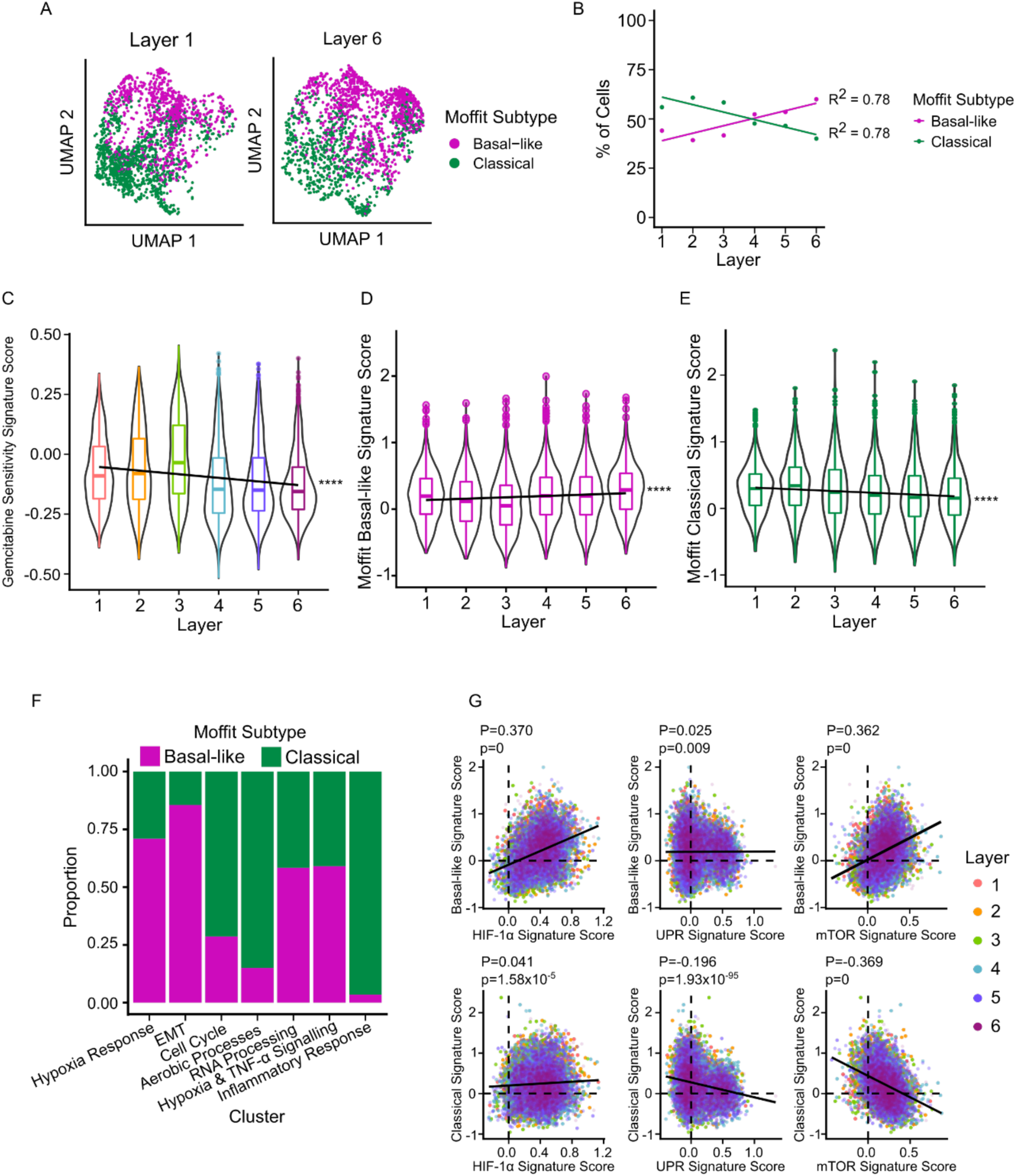
Microenvironmental gradients contribute to an increase in the proportion of Moffit basal-like organoid cells. (A) UMAP plot showing the distribution of organoid cells classified by their maximal Moffit signature scores in Layer 1 (outer) and Layer 6 (inner) of TRACER. (B) Quantification of organoid cell subtype classification proportions by layer showed a graded increase in the proportion of basal-like organoid cells towards the inner layers of TRACER. (C) Tiriac et al. PDO-derived gemcitabine sensitivity score for organoid cells in each layer of TRACER^59^. Tumour cell intrinsic changes in predicted sensitivity were observed across TRACER layers, consistent with the observed increasing proportion of basal-like organoid cells in the inner layers of TRACER. (D) The Moffit basal-like signature score exhibited a significant graded increase across TRACER layers (slope of the best fit line vs. 0, ****p< 0.0001) (E) The Moffit classical signature score exhibited a significant graded decrease across TRACER layers (slope of the best fit line vs. 0, ****p<0.0001). (F) Quantification of subtype proportion for each organoid cell cluster revealed an association between the hypoxia response-related clusters and more basal-like organoid cells, in addition to an expected association between the EMT cluster and more basal-like organoid cells. (G) Assessment of the correlation between the basal-like and classical gene signature scores and known hypoxia response pathway gene signature scores for all organoid cells retrieved from TRACER culture (HIF-1α Transcription Factor Network – Pathway Interaction Database, Unfolded Protein Response (UPR) - Reactome and mTOR_4Pathway – Pathway Interaction Database) implicated each hypoxia response pathways to different degrees in promoting a more basal-like organoid cell phenotype (P: Spearman correlation coefficient).

We next examined whether the observed increase in the proportion of basal-like organoid cells was driven by an increase in the expression of the basal-like gene signature or a decrease in the expression of the classical gene signature. We observed both a significant graded increase in the Moffit basal-like signature score and a significant graded decrease in the Moffit classical gene signature scores from Layer 1 (outer) to Layer 6 (inner) (slope of the best line vs. 0, p < 0.0001; **Figs. 3D-E, Table S2**). Furthermore, our analysis showed no significant difference in the magnitude of these gradients, suggesting that both an increase in basal-like gene expression and a decrease in classical gene expression contributed to the observed increase in the proportion of basal-like organoid cells from the outer to inner layers. Reinforcing this result, increasing trends were also observed in the expression of EMT and TGF-β signalling gene sets, previously associated with the basal-like subtype, and a decreasing trend was observed in the expression of a pancreatic progenitor gene set, which has previously been associated with the classical subtype (**Fig. S4B-D, Table S2**)^3,55^. Taken together, this data supports the idea that microenvironmental gradients contribute to the determination of transcriptional subtypes in PDAC.

Given the predictable variation in organoid cell transcriptional subtype and hypoxia response signatures across the TRACER layers, we next assessed the proportion of basal-like organoid cells in each of the single cell clusters associated with hypoxia. We observed that the majority of cells in the hypoxia response and hypoxia response/TNF-α signalling clusters were classified as basal-like (**Fig. 3E**). We also assessed the correlation between three known hypoxia response pathways and the transcriptional subtypes to explore whether specific hypoxia response mechanisms may promote the basal-like phenotype (**Fig. 3F, Table S2**)^56^. In this analysis, we found that the Moffit basal-like gene signature score was positively correlated (Spearman correlation coefficient *ρ* = 0.370, p = 0) with the HIF-1α transcription network signature score, whereas the classical gene signature score was very weakly correlated with this pathway (*ρ* = 0.041, p = 1.58×10^-5^). Conversely, the Moffit classical signature score was negatively correlated with the unfolded protein response (UPR) signature score (*ρ* = -0.196, p = 1.93×10^-95^), whereas the basal-like signature score was weakly correlated with this pathway (*ρ* = 0.025, p = 0.009). Finally, we examined the correlation with mTOR signalling as the AKT/mTOR pathway has been implicated in facilitating the translation of HIF-1α in PDAC^57^ and found that the mTOR signalling signature score was positively correlated with the Moffit basal-like signature score (*ρ* = 0.362, p = 0) and negatively correlated with the classical signature score (*ρ* = -0.369, p = 0). This correlation is consistent with previous work showing enrichment of mTOR signalling in squamous/basal-like PDAC tumour cells^58–60^. Additionally, work in an intraductal organoid transplantation model of PDAC has shown that both mTOR signalling genes and basal-like signature genes are enriched in organoids that rapidly progress to an invasive phenotype^61^. Collectively, our analysis suggests that the cellular response to hypoxia is strongly associated with the increase in basal-like organoid cells across the layers of TRACER, with multiple hypoxia response pathways likely contributing to this shift in organoid cell state.

### Hypoxia is sufficient to drive the enrichment of a more basal-like phenotype in PDAC organoid cells

Having identified a clear association between hypoxia response and the increased proportion of basal-like organoid cells in the inner layers of TRACER in our scRNA-seq data, we next sought to determine whether hypoxia plays a causal role in driving the enrichment of a basal-like phenotype. We first confirmed the enrichment of basal-like gene expression in the inner layers of TRACER by performing qPCR for selected markers of the classical (GATA6, LYZ) and basal-like (KRT5, S100A2) subtypes identified in previous studies (**Fig. S7A**)^3,7,12^. Next, to isolate the effect of hypoxia on the expression of these markers, we compared the expression of these genes in organoid cells cultured in a 0.2% O_2_ hypoxia chamber for 24 h to organoids cultured in normoxia using qPCR. We also quantified expression of these markers in Layer 1 (outer) and Layer 6 (inner) of TRACER (the extremes of the oxygen and microenvironmental gradients) relative to organoids cultured in normoxia for reference (**Fig. 4A, Table S2**). Consistent with our scRNA-seq analysis, we observed a significant decrease in the expression of the classical marker genes GATA6 and LYZ in hypoxia (GATA6: p < 0.05; LYZ: p < 0.0001) and a significant increase in the expression of the basal-like marker gene KRT5 (p < 0.05) in hypoxia. Furthermore, the changes in expression levels of these genes were similar to those observed in the hypoxic inner layers of TRACER. These data are consistent with a causal role for hypoxia in driving organoid cells towards a more basal-like transcriptional phenotype, that predominates over the other microenvironmental gradients present in TRACER. Interestingly, there was no significant change in the expression of basal-like marker S100A2 in 0.2% O_2_; however, there was a significant increase in S100A2 expression in Layer 6 (inner) of TRACER (p < 0.05), suggesting that while hypoxia plays a dominant role in the observed reprogramming, other microenvironmental gradients may also contribute to the underlying changes in gene expression observed in organoid cells cultured in TRACER.

**Figure 4:**
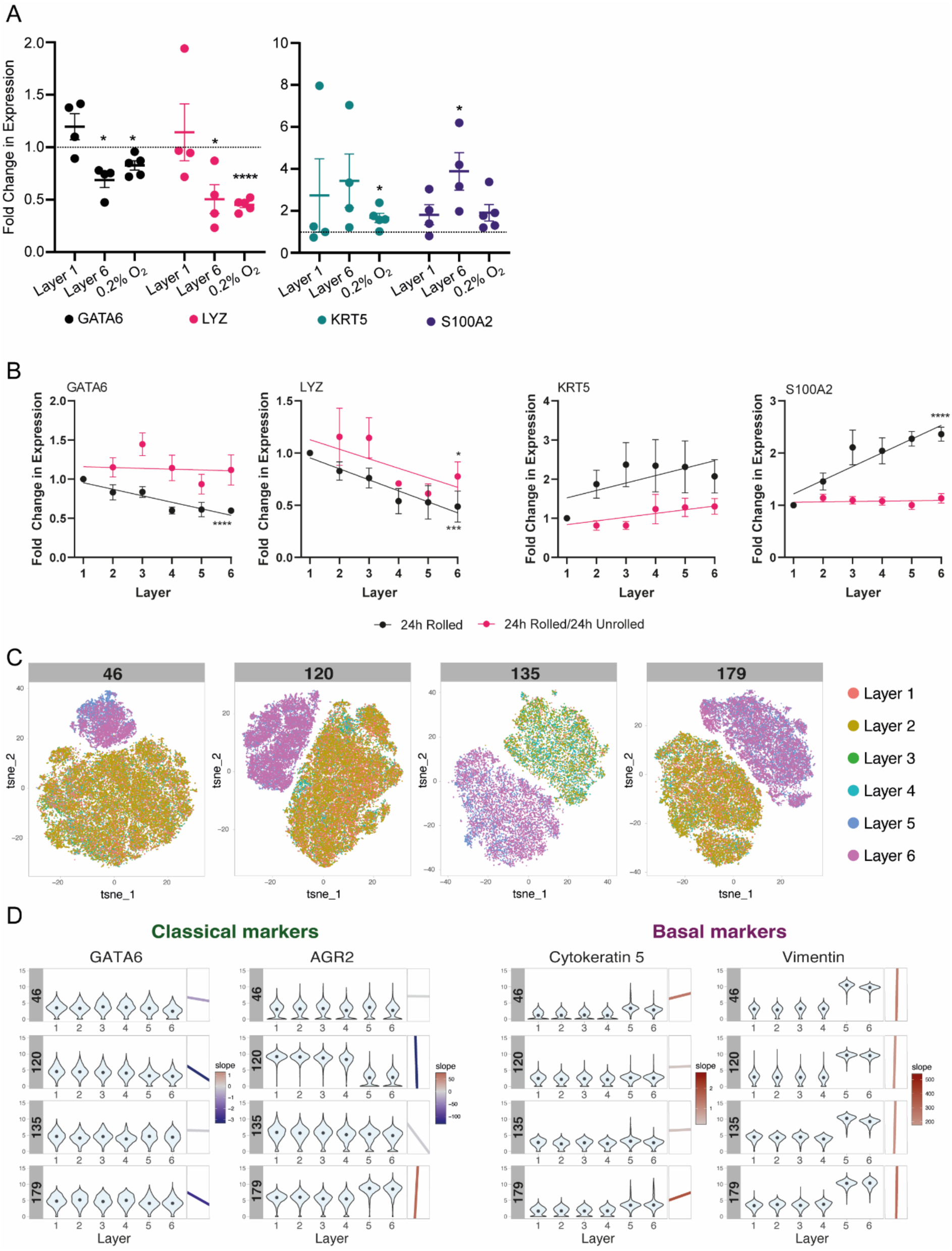
Hypoxia causes changes in the expression of transcriptional subtype markers in multiple PDAC organoid lines, which may be partially reversible. (A) Comparison of the gene expression of GATA6, LYZ (classical) and KRT5, S100A2 (basal-like) markers in Layer 1 (outer) and Layer 6 (inner) of TRACER and in 0.2% pO_2_ relative to organoid cells cultured in normoxia after 24 h showed decreases in classical gene expression and an increase in KRT5 expression in hypoxia, suggesting hypoxia plays an important role in promoting a more basal-like organoid cell phenotype (TRACER: 4 independent experiments with n=3 TRACERs, 0.2% pO_2_: 5 independent experiments with n=3 technical replicates; t-test vs. 1, *p < 0.05, ****p < 0.0001; mean ± SEM). (B) Comparison of the expression of classical and basal-like marker genes between organoid cells cultured in rolled TRACERs for 24h and organoid cells cultured in rolled TRACERs for 24h followed by an additional 24h of culture in an unrolled configuration (normoxia). Graded changes in GATA6 and S100A2 expression were abrogated after additional 24h of culture in normoxia, suggesting a specific role for hypoxia in transcriptional reprogramming towards a basal-like phenotype and potential reversibility when returned to normoxia (slope of the best fit line vs. 0, ****p < 0.0001; 24h rolled: 4 independent experiments with n=3 TRACERs, 24h rolled/24h unrolled: 3 independent experiments with n=3 TRACERs; gene expression normalized to Layer 1 (outer), mean ± SEM). (C) T-SNE visualisation of high dimensional analysis performed using a panel of PDAC-specific proteins markers by CyTOF in different organoids models (PPTO.46, 120, 135, 179). Cells are colour-coded based on their TRACER layer of origin. (D) Analysis of the expression of GATA6, ARG2 (classical) classical and CK5, Vimentin (basal-like) in TRACER seeded with different organoid models (organoid model number indicated in grey). Violin plots summarize the metal intensity of each cell and black dots indicate the mean expression per layer for each organoid model and protein marker. The insets on the right side of the violin plots indicates the slope of the best fit line, with lines colour-coded by slope values and sign (blue: negative slope; red: positive slope).

We also explored the reversibility of the observed microenvironment-dependent reprogramming of the organoid cells towards a basal-like state. To this end, we assessed the effect of culturing cells in the rolled configuration of TRACER for 24 h followed by unrolling the construct and allowing an additional 24 h of culture in a now normoxic environment. After the additional 24 h of culture in the non-rolled configuration, we observed that the graded expression of GATA6 expression across the TRACER layers was lost (**Fig. 4B, Table S2**). Similarly, the significantly increasing gradient in S100A2 expression (basal-like gene) observed across the TRACER layers was abrogated after the additional 24 h of culture in a, now, normoxic environment. However, we did not observe any significant changes in the graded expression of LYZ (classical) or in the expression of KRT5 (basal-like). Using cytometry time of flight (CyTOF) to assess expression of subtype markers at the protein level, we observed similar trends in the expression of classical markers GATA6 and AGR2^59^; however, no significant changes were observed for basal-like markers Vimentin (VIM)^63^ and Cytokeratin 5 (CK5)^12^ after reoxygenation (**Figure S7B**). Given the robust association between GATA6 expression and the classical subtype^12^, our observations provide more evidence of a causal relationship between oxygen level and transcriptional subtype. Furthermore, these observations suggest that the transcriptional subtypes are plastic and that the increase in basal-like organoid cells observed in hypoxia may be at least partially reversible if normoxia is restored.

We then assessed the relationship between hypoxia and transcription sub-type in a larger cohort of organoid lines. To do this, we leveraged a CyTOF barcoding strategy that allowed the simultaneous measurement of multiple protein of interest in a large number of samples. We selected various PDAC-relevant protein markers with validated CyTOF antibodies available for our analysis, including PDAC subtype markers (SI Table S5). Using this strategy, we tested TRACERs generated from 4 organoid lines (PPTO.46, 120, 179, and 135). TRACERs seeded with each line were cultured for 4 days then rolled for 24h, as in the scRNA-seq analysis, to allow the generation of an oxygen gradient across the layers. TRACERs were then unrolled for analysis of the cells from each layer using CyTOF. The presence of an oxygen gradient was verified by the increased uptake of the Telox hypoxia probe^64^ in the inner layers compared to outer layers (**Figure S7C**). High dimensional analysis using all measured protein markers for each organoid model indicated that organoid cells from layers 1 to 4 showed distinct expression profiles from organoid cells isolated from layers 5 and 6 as shown by TSNE plots (**Figure 4C**) correlating with the more severe hypoxia in layers 5 and 6 (**Figure S7C**). We then assessed the extent to which the expression of each marker was graded from layer 1 to 6 (**Figure S7D**). Examining the expression of basal-like and classical markers specifically, we observed graded trends consistent with our analysis in PPTO.46. Specifically, the expression of classical markers GATA6 and AGR2 decreased from the outer normoxic layers to the inner hypoxic layers in all organoid models tested, with the exception of AGR2 in the PPTO179 organoid model which showed no gradient (**Figure 4D**). Further, the expression of basal-like markers CK5 and VIM increased in the hypoxic inner layers in all organoid models tested (**Figure 4D**). Collectively, these results reinforce the findings from our scRNA-seq analysis of TRACER fabricated with the PPTO46 organoid model, namely that hypoxic gradients play a contributing role in determining the transcriptional subtype of PDAC tumour cells.

## Discussion

The interaction between tumour cells and the TME undoubtedly plays an important role in supporting tumour growth, progression, and response to treatment^15,16,18^. However, experimental limitations including difficulty accessing tumour samples with sufficient cellularity for analysis and a lack of disease-representative *in vitro* models have hindered explorations of the specific impacts of microenvironmental features on transcriptional phenotypes in PDAC^59,65,66^. Although scRNA-seq has offered deeper insight into the transcriptional heterogeneity of tumour cells, it has not been possible to correlate transcriptional phenotypes with the tumour cell-extrinsic features of a defined location in the TME. To address this challenge, we applied scRNA-seq to PDAC organoid cells cultured in TRACER, an engineered paper-based 3D *in vitro* model that models disease-relevant microenvironmental gradients, to perform a spatially resolved, single-cell analysis of the relationship between tumour cell phenotypes and microenvironmental features.

In this proof-of-concept study, we first assessed the extent of transcriptional heterogeneity in TRACER and evaluated how this heterogeneity varied between the TRACER layers, which correspond to different levels of hypoxia. Our analysis identified 7 transcriptional organoid cell phenotypes that were present in each layer but in varying proportions. These clusters exhibited enrichment of gene sets related to hypoxia response, EMT, cell proliferation, aerobic metabolism, RNA processing, TNF-α signalling, and IFN-response, in line with gene expression patterns identified in previous work^14,22,36,67–69^. Notably, the representation of all clusters in each layer suggests that TRACER supports transcriptional heterogeneity in organoid cells consistent with the level of heterogeneity observed in previous single cell transcriptomic studies of PDAC tumours and PDAC PDOs^14,36,67^. Importantly, the proportion of cells in each cluster differed in each layer, with an increase in the proportion of hypoxia-response related clusters and a decrease in the proportion of the cell cycle and aerobic processes clusters in the inner layers of TRACER, consistent with the expected response to hypoxia^32^. Accordingly, the transcriptional patterns observed in this study reflect the organoid cell response to hypoxia, which is a critical aspect of PDAC tumours *in vivo* that has largely been overlooked in studies of PDOs *in vitro* to date^18^. Furthermore, as all organoid cells were originally derived from the same PDO source, and our previous work has shown minimal organoid cell death in TRACER after 24h in rolled culture^32^, we propose that changes in cluster proportion are likely due to microenvironment-related transcriptional reprogramming rather than the enrichment of specific subpopulations in the inner layers of TRACER. We also noted that some cluster proportion trends were more robustly associated with the microenvironmental gradient in TRACER than others, which suggests that not all subpopulations of cells are equally sensitive to microenvironmental conditions; further exploration of whether there are subpopulations of tumour cells that are relatively microenvironment-insensitive will be an interesting area for future work. In short, the microenvironmental gradient present in TRACER offers an important additional dimension for understanding factors that affect transcriptional heterogeneity in PDAC organoid cells.

In parallel, we also identified gene sets enriched based on gene expression correlation with TRACER layer number, suggesting that microenvironmental conditions had a particularly significant impact on the expression patterns of these gene sets. Here, we found that gene sets with the most pronounced trend in enrichment across the layers were related to hypoxia and stress response; gene sets related to metabolism, cell proliferation, major signalling pathways, EMT and protein localization also showed enrichment correlated with TRACER layer number. Though many of these gene sets showed enrichment consistent with our expectations and/or have been identified as being enriched in PDAC tumour samples or organoids^14,22,67–69^, our findings provide additional and direct evidence that enrichment of these gene sets is promoted by microenvironmental conditions^21,41,42,70^. Importantly, as only organoid cells were present in the TRACER construct, these transcriptional changes resulted from the direct effect of microenvironmental gradients on organoid cells rather than being mediated by other cell populations, such as stromal cells. This analysis offers several gene sets that may be interesting targets for future investigations. Specifically, based on our previous work^70^ and the correlation between metabolism-related gene sets and TRACER layer number observed in this study, we anticipate microenvironmental gradients of key metabolites, such as lactate and fatty acids, are likely present across the TRACER layers. Thus, further explorations of the metabolic adaptations of PDAC organoid cells in TRACER will be a valuable avenue for future study, as tumour metabolism can significantly impact anti-tumour immune responses^71–73^ and therapeutic efficacy^74–76^. Furthermore, the unique design of TRACER offers a valuable platform for such studies as it enables spatially resolved metabolomic analysis not achievable using other experimental approaches^32,33^. Such future investigations of the downstream effects of microenvironment-related changes in tumour cell phenotypes in TRACER will provide an opportunity to gain a more comprehensive understanding of the interactions between tumour cell phenotypes and microenvironmental conditions.

As the basal-like and classical transcriptional subtypes have prognostic value in PDAC, we focused here on examining the effect of microenvironmental gradients on transcriptional subtype identity and the expression of subtype marker genes^3,6,7^. Our results showed layer-specific changes in the proportion of basal-like and classical organoid cells, suggesting that microenvironmental gradients contribute to the determination of the basal-like and classical transcriptional subtypes. Specifically, we identified hypoxia as a dominant microenvironmental factor that promotes a more basal-like phenotype, which aligns with previous work showing an association between hypoxia response gene expression and the basal-like phenotype^10,12,60,77^. Importantly, by using TRACER, which has experimentally confirmed hypoxia in the inner layers, in conjunction with a hypoxia chamber, our work mitigates the limitations associated with inferring microenvironmental conditions from gene expression to confirm a role for hypoxia in subtype determination^10,32^. Although the precise mechanism by which hypoxia promotes a more basal-like phenotype remains to be determined, our finding that the basal-like and classical transcriptional signatures were most strongly correlated with different hypoxia response pathways suggests multiple pathways contribute to promoting a more basal-like phenotype. Notably, we identified the mTOR pathway as being significantly correlated with the basal-like signature score and significantly negatively correlated with the classical signature score, in line with previous work showing enrichment of mTOR signalling in basal-like PDAC tumour cells^58–60^. Given the interest in mTOR as a therapeutic target in PDAC, further investigations of the relationship between mTOR signalling and the transcriptional subtypes may be of significant interest^78,79^. However, it should be noted that while we used organoid cells from three additional patients to validate our initial findings, emerging evidence suggests that most organoid models reflect the classical subtype at baseline^6^. Thus, future work should assess how transcriptional heterogeneity in the initial sample impacts microenvironment-dependent reprogramming using a larger cohort of organoid models. Taken together, our findings do however add to growing evidence that the basal-like subtype is associated with certain microenvironmental features^6,21,80,81^, and that the TME contributes to the determination of transcriptional subtypes in PDAC.

Given that basal-like tumours are associated with a more aggressive and treatment resistant phenotype^7,10^, the shift towards a more basal-like phenotype in hypoxia may have implications for response to therapy. In line with the observed increase in basal-like organoid cells in the inner layers of TRACER, our previous work has shown that organoid cells in the inner layers of TRACER are less sensitive to treatment with gemcitabine chemotherapy^32^. However, it remains unclear whether microenvironment-dependent reprogramming to a more basal-like phenotype can be reversed. In this study, we demonstrated that organoid cells removed from the inner layers of TRACER and cultured for an additional 24 h in normoxia showed abrogation of the GATA6 and S100A2 expression gradients observed in hypoxia, suggesting the potential for at least partial reversibility of hypoxia-induced reprogramming. Accordingly, further exploration of whether oxygenation can restore cells to a classical phenotype and, if so, whether this could improve chemosensitivity in PDAC tumour cells would be a valuable area for future study. In any case, it is clear from our study that microenvironmental conditions, such as hypoxia, significantly impact organoid cell phenotypes and therefore should be incorporated in organoid-based studies of PDAC biology and response to therapy. Ultimately, considering these key tumour-microenvironment interactions could pave the way to the identification of new treatment strategies that more comprehensively target the complex factors that contribute to tumour growth and progression in PDAC.

## Conclusion

In this study, we harnessed the TRACER platform in conjunction with PDAC patient-derived organoids to perform a spatially resolved, single cell transcriptomic analysis that enabled identification of distinct cell phenotypic clusters, the relative proportions of which varied with microenvironmental conditions. The precise control over cellular components and established microenvironmental gradients of oxygen and other small molecules in TRACER offered important advantages over conventional sequencing analysis of tumour samples and organoids. Significantly, we found that hypoxia is a key microenvironmental player in determining transcriptional subtypes and likely has a causal role in driving cells towards a more basal-like phenotype. We anticipate that this work will provide foundational understanding for further investigations into the mechanisms contributing to microenvironment-dependent transcriptional reprogramming, particularly with regards to the major transcriptional subtypes in PDAC. Elucidating these mechanisms may facilitate the identification of therapeutic targets that acknowledge the importance of the TME in influencing tumour cell phenotypes.

## Methods

### Patient-Derived Organoid (PDO) Culture

Experiments were conducted with established pancreatic ductal adenocarcinoma tumour organoids (PMLB PPTO.46 and PMLB PPTO.120) from the Princess Margaret Living Biobank at the University Health Network, Ontario, Canada. Approval for use of the materials was provided by the University of Toronto Research Ethics Board (protocol number 36107). PPTO.46 and PPTO.120 cells were maintained in Advanced DMEM/F-12 (Gibco) supplemented with 2 mM GlutaMax, 10 mM HEPES (Gibco), 1% penicillin/streptomycin, 1X W21 supplement (Wisent), 1.25 mM N-acetyl-L-cysteine, 10 nM Gastrin I (1-14), 10 mM nicotinamide (Sigma-Aldrich), 50 ng/mL recombinant human EGF (Gibco), 100 ng/mL recombinant human noggin, 100 ng/mL recombinant human FGF-10 (Peprotech), 0.5 μM A83-01 (Tocris Biosciences), 10 μM Y-27632 (Selleck Chemicals), 20% v/v Wnt-3a conditioned media, and 30% v/v human R-spondin1 conditioned media (Princess Margaret Living Biobank, Toronto, Canada). Media for PPTO.120 cells was further supplemented with 2.5 μM CHIR99021 (Tocris Small Molecules). PPTO.46 and PPTO.120 cells were cultured in 48-well polystyrene plates in 50 uL domes of Growth Fator Reduced Phenol Red-Free Matrigel Matrix (Corning Life Sciences) with 500 uL of complete media. Media was refreshed twice per week and cells were passaged once per week (PPTO.46 1:8 split ratio, PPTO.120 1:4 split ratio). All cultures were maintained in a humidified atmosphere at 37°C and 5% CO_2_. Organoid models were used to a maximum of passage 30.

### TRACER System Fabrication

TRACER scaffolds and seeding devices were fabricated as previously described^32^. Briefly, TRACER scaffolds were manufactured by infiltrating cellulose paper strips (0.5 x 12 cm, Miniminit Products) with poly(methyl methacrylate) (PMMA, 120 kDa) dissolved in acetone (Sigma-Aldrich) at 0.175 g/mL with an Allevi Beta 1 3D printer. Seeding devices were fabricated with poly(dimethyl siloxane) PDMS Sylgard 184 (Dow Silicones Corporation) using a monomer to cross-linker ratio of 9:1 (w/w) for the device lids and 20:1 (w/w) for the base. Nylon tape strips with a thickness of ∼100 μm were used to mould grooves in the PDMS base for positioning of TRACER scaffolds as previously described^34^. A 1mm biopsy punch (Integra Life Sciences) was used to create inlet and outlet holes in the PDMS lids aligned with each of the six regions for cell infiltration into the TRACER scaffold. The device components were sterilized by UV light exposure and assembled on a polycarbonate base for easier handling, with the TRACER scaffolds sandwiched between the grooves cast in the base and the PDMS lids.

### TRACER Seeding

TRACER strips were seeded with organoid cells as previously described^32^. Briefly, organoids were retrieved from the Matrigel domes and dissociated to single cells with TrypLE (Gibco). Organoid cells were then collected, pelleted, and re-suspended in a 25:75 (v/v) mixture of neutralized Type 1 bovine collagen (PurCol 3 mg/mL; Advanced Biomatrix) and Growth factor reduced phenol red-free Matrigel (Corning Life Sciences) at a density of 10 x 10^6^ cells/mL. Collagen was prepared by mixing 500 uL of collagen with 62.5 uL of 10x minimum essential medium (Life Technologies) and neutralized with 0.8 M NaHCO_3_ to reach a neutral pH. ECM blends were kept on ice after preparation and used within 10 mins for seeding of TRACER strips. 5 uL of cell-gel suspension was then injected through the central inlet hole in the PDMS seeding device corresponding to each layer of TRACER. Finally, the seeding device was placed in a humidification chamber at 37 °C for 45 mins to allow for gelation.

### TRACER Experimental Approach

After gelation, the cell-infiltrated TRACER strips were transferred a 12 well plate with 3 mL of complete organoid growth media for 4 days. After 4 days in unrolled culture, TRACER scaffolds were rolled around a custom aluminum mandrel and the media volume was topped up to 4 mL with complete organoid media. TRACER scaffolds were cultured in the rolled configuration for 24 h. For unrolled controls, TRACER strips were sectioned into individual layers and cultured in either normoxia or a 0.2% pO_2_ hypoxia chamber for 24 h in parallel with the rolled constructs, as applicable.

Organoid cells were retrieved from the scaffold at the completion of the experiment by unrolling the TRACER scaffolds and sectioning the strip into individual layers. The individual layers were then incubated in a solution of HBSS supplemented trypsin 0.5x for 30 mins on an orbital shaker at 900 rpm and 37°C^34^. The samples were then agitated by repeated pipetting to further dislodge the cells, and Advanced DMEM/F-12 media supplemented with 10% FBS was used to neutralize the digestion reaction. Cells were then pelleted by centrifugation and processed for downstream analysis.

### 10x Single Cell Sample Preparation

Three TRACER scaffolds were prepared as described above with PPTO.46 ODCs. After 24 h in the rolled configuration, organoid cells were retrieved from the scaffold and counted. The scaffold with the greatest number of cells per layer on average was sent for sequencing. Samples were prepared for sequencing as outlined by 10x genomics Single Cell 5’ v2 Reagent Kits user guide. Cells were washed and counted and viability assessed using haemocytometer (ThermoFisher). Following counting, the appropriate volume for each sample was calculated for a target capture of 2000 cells and loaded onto 10X single cell chip K. After droplet generation, samples were transferred onto a pre-chilled 96 well plate (Eppendorf), heat sealed and incubated overnight in a Veriti 96-well thermocycler (ThermoFisher). The next day, sample cDNA was recovered using Recovery Agent provided by 10x and subsequently cleaned up using a Silane DynaBead (ThermoFisher) mix as outlined by the user guide. Purified cDNA was amplified for 16 cycles before being cleaned up using SPRIselect beads (Beckman). Samples were run neat on a Bioanalyzer (Agilent Technologies) to determine cDNA concentration. 5’ v2 cDNA libraries were prepared as outlined by the Single Cell 5’v2 Reagent Kits user guide with modifications to the PCR cycles based on the calculated cDNA input.

### Single-cell RNA Sequencing

The library size was determined using Agilent High Sensitivity DNA Kit on the Bioanalyzer instrument (Agilent Technologies). The sample was then pooled with other 10X libraries, quantified using the Qubit dsDNA HS Assay Kit on the Qubit 2.0 Fluorometer (Invitrogen), and subsequently normalized to 4 nM using elution buffer (Qiagen) with 0.1% Tween20 (Sigma). The 4 nm pool was denatured using 0.2N NaOH at equal volume for 5 minutes at room temperature. Library pool was further diluted to 20 pM using HT-1 (Illumina) before being diluted to a final loading concentration of 12.5 pM. This multiplexed pool was sequenced with the following parameters on the MiSeq platform with a Micro flowcell (Illumina): Read 1 – 28 cycles, Read 2 – 91 cycles, Index 1 – 8 cycles. The number of actual cells captured was estimated based on this shallow sequencing data analyzed using the CellRanger pipeline (10X Genomics). The target number of reads for the sample were adjusted based on the sequencing results of the MiSeq pool. The sample was then re-pooled with other 10X libraries, quantified using the Qubit dsDNA HS Assay Kit on the Qubit 2.0 Fluorometer (Invitrogen), and subsequently normalized to 1.5 nM using Low TE buffer (Invitrogen). The 1.5 nm pool was denatured using 0.2N NaOH according to Illumina guidelines for 8 minutes at room temperature. Library pool was further diluted to 350 pM using 400mM Tris-HCl. This multiplexed pool was sequenced with the following parameters with an S2 flow cell on the NovaSeq 6000 platform (Illumina): Read 1 – 28 cycles, Read 2 – 91 cycles, Index 1 – 8 cycles.

### Single-cell RNA Sequencing - Data Processing and Clustering

Demultiplexing, alignment to reference genome GRCh38, and quantification, was performed from FASTQ using Cell Ranger 6.0. For each TRACER layer, cell libraries were filtered out if they contained transcripts from less than 500 genes, or those with a proportion of mitochondrial transcripts greater than four median absolute deviations over the median. Cell libraries from each TRACER layer were normalized using the pool and deconvolute approach available in the *scran* R package^82^. Normalized gene abundances per layer were then combined into a single gene by cell dataset. After assessing the relative abundance of housekeeping genes between layers (**Figure S8A-B**), no further normalization was necessary.

Clustering and 2D representation of cell libraries in gene expression space were performed after reducing the feature space to its most informative components in a two-step process. First, highly-variable genes (HVGs) were identified, then principal component analysis was used to summarize these gene expression vectors into a smaller set of orthogonal and informative components. UMAP was used to summarize these multi-dimensional component vectors representing each cell into 2D for visualization^83^. Clustering of cells was performed on a pruned k-nearest neighbour (kNN) graph as implemented in the *Seurat* R package (v4.1.0)^84^. The optimal number of clusters was determined by assessing each clustering’s robustness to changes in input parameters - specifically number of nearest neighbours in the kNN graph, the resolution parameter used to cluster that graph, and the method used to determine HVGs. HVGs were assigned as either the thousand genes with the greatest variance relative to their mean expression, those genes with significantly more variance than expected given their mean expression (FDR < 0.1), determined by DUBSTEP, or all genes^85,86^. Robustness was assessed as the Adjusted Rand Index (ARI) for each pair of cluster solutions resulting in the same number of clusters. Since changing the input genes (HVGs) caused the most difference in clustering solutions, specific priority was given to the solution that was robust to HVG selection. Optimal clustering solutions contained either 2 or 7 clusters, the latter clusters being a subset of the former. Accordingly, we proceeded with analysis of 7 clusters to better assess transcriptional heterogeneity within the TRACER construct.

### Single-cell RNA Sequencing – Data Analysis

Differential gene expression was calculated by Wilcoxon rank-sum test using the *scClustViz* R package^87^. Gene Set Enrichment Analysis was performed using a custom gene set collection designed for Enrichment Map containing both GO biological process terms as well as a compilation of pathway databases available at ref. 67^88–90^. Clusters were annotated based on their enrichment of related gene sets sourced from multiple databases and the maximum normalized enrichment score (NES) for these gene sets was reported where applicable.

The hypoxia score was calculated as described in Connor et al.^83^. Briefly, the hypoxia score is calculated as the difference in median expression of 20 genes with the highest correlation and 10 genes with the lowest correlation with stratifying tumours by hypoxia status. Other signature scores were calculated using the AddModuleScore function in Seurat. For the subtype scoring and determination, the Moffit subtype scores were determined using the 25 marker genes for each subtype^84^, with the combined Moffit score calculated as max(0, classical score)-max(0, basal-like score)^14^. The Chan-Seng-Yue subtype scores were determined using the top 100 ranked genes for the classical and basal-like signatures^87^, and the Raghavan subtype scores were determined using the top 30 genes per subtype^6^. The gemcitabine sensitivity signature used was the refined PDO gemcitabine sensitivity signature from Tiriac et al^26^. The EMT signature was sourced from a consensus gene set^91^. The pancreatic progenitor gene signature was sourced from Balli et al^55^. All other gene sets were sourced as indicated from the compiled pathway database used for GSEA.

Data visualization was performed using Seurat v4.1.0, scCustomize, and ggplot2^84,92,93^.

### Gene Expression Analysis by qPCR

TRACERs were unrolled following 24 h of rolled culture and sectioned into individual layers. The RNeasy Mini Kit (Qiagen) was used to isolate RNA from ODCs in each layer. Individual layers and unrolled controls were washed briefly in PBS and submerged in 500 uL RLT Lysis Buffer with 5 μL β-mercaptoethanol in 1.5 mL centrifuge tubes. Tubes were vortexed at room temperature for 10 mins. The remainder of the RNA extraction was performed in accordance with the manufacturer’s instructions. The concentration of extracted RNA was measured using a NanoDrop spectrophotometer (ThermoScientific). 500 ng RNA was converted to cDNA by reverse transcriptase PCR with qScript cDNA SuperMix (QuantaBio). 7 ng of cDNA was used for qPCR and amplified with gene-specific primers (Table S4, MilliporeSigma) and the PowerSybr Green Master Mix (Applied Biosystems). The qPCR was performed with a CFX-384 thermocycler (BioRad), and data were acquired using Bio-Rad CFX Manager 3.1. Gene expression was normalized to a housekeeping gene (RPL13a) and the 2^-ΔΔCt^ method was used to determine relative expression. Data was normalized to Layer 1 (outer) of each TRACER2 construct (excluding Fig. 4A) or to the unrolled normoxic control (Fig. 4A).

### Sample Preparation for Mass Cytometry

TRACER samples were prepared for analysis with CyToF as previously described^34^. Briefly, TRACER rolls and unrolled control samples were incubated at 37 °C in complete organoid media with 50 μM of 5-Iodo-2’deoxyuridine (IdU, Fisher Scientific) and 20 uM of Telox^64^ (courtesy of Mark Nitz) for 6h prior to unrolling. After unrolling, organoid cells were retrieved from TRACER after 24 h in rolled culture and from hypoxic and normoxic control samples as described above. Cells were pelleted by centrifugation at 300g for 5 minutes, washed once with PBS and then fixed with PFA 4% for 10 min at room temperature. PFA was then removed, and cells were washed twice with PBS. Individual samples were then transferred to a V-bottom 96 well plate for barcoding. To barcode samples, we used a barcoding strategy combining m-DOTA barcoding and the Cell-ID 20-Plex Pd Barcoding Kit (Standard BioTools, San Francisco, USA). Briefly, each tube of the Cell-ID 20 Plex Barcoding kit containing 3 premixed palladium isotope was mixed with one m-DOTA (Y89, Ce140, Tb159, Ho165) to create a barcoding scheme allowing the unique barcoding of all the samples. The barcoding scheme is presented in **Figure S9A**. After barcoding, all samples were pooled in a single tube. All cells were then permeabilized using saponin 0.2% for 30 min at room temperature. After permeabilization, cells were stained with a cocktail of metal-conjugated antibodies (details of the antibodies presented in supplementary **Table S5**) and diluted in cell staining buffer (Standard BioTools, San Francisco, USA). After staining, cells were washed and DNA was stained using 125 nM of Cell-ID Intercalator Ir (Standard BioTools) overnight at 4°C. The next day, cells were washed twice with cell staining buffer (Standard BioTools) and then washed twice with Cell Acquisition Solution Plus (Standard BioTools). Pellets were then resuspended in Cell Acquisition Solution Plus containing EQ six element calibration beads (Standard BioTools) and analyzed using CyTOF XT (Standard BioTools). Event rate during acquisition was maintained between 200 and 300 events per second.

### Analysis of Mass Cytometry Data

After acquisition, normalization and randomization were applied using the CyTOF Software algorithm (Standard BioTools). FCS files were then debarcoded by first isolating populations based on the m-DOTA barcodes (**Figure S9B**). Files containing only cells with a unique m-DOTA barcode were then further debarcoded using the CyTOF Software for debarcoding based on palladium isotopes. After debarcoding, each sample was saved as a *.fcs file for downstream analysis. Single cell populations were gated using Gaussian parameter discrimination as previously described^94^. Results are presented as violin plots that summarize single cell-level data, with dots that represent the median metal intensity of each layer in each TRACER. For each marker in each organoid model, the slope of the linear regression was calculated and depicted by a line colour-coded by the slope. T-SNE plots were also generated using all phenotypic markers (**Table S5**) using R.

### Statistical Analysis

Statistical analysis was performed using RStudio with R version 4.1.2 and GraphPad Prism 9.0.1 (GraphPad Software). p < 0.05 was considered statistically significant. Power calculations were not used to determine the number of replicates for each experiment. The details of the statistical analyses used are indicated in the figure captions and summarized in Table S2.

## Acknowledgements

This work was funded by a Vanier Canada Graduate Scholarship to NLB, a PRiME fellowship to BTI, a fellowship from CFREF Medicine by Design to SL, an Ontario Graduate Scholarship to JLC, a Canadian Institute of Health Research (CIHR) Project Grant (PJT-17528) to APM.

## Data availability statement

Data will be provided upon reasonable request to the authors.

