## Supplementary information for "Single-cell analysis of an engineered organoid-based model of pancreatic cancer identifies hypoxia as a contributing factor in the determination of transcriptional subtypes"

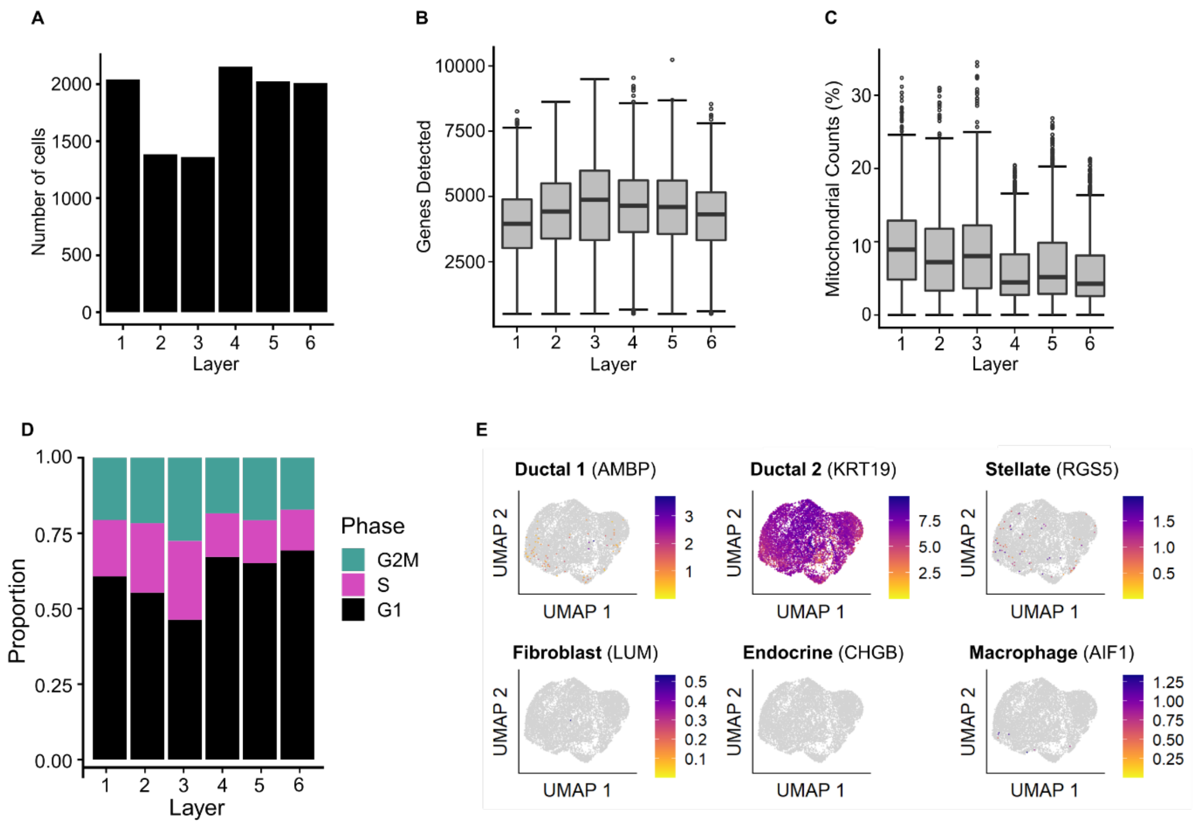


***Supplementary Figure S1: Quality control of scRNA-seq of TRACER fabricated with PPTO.46 PDAC organoid cells after pre-processing.*** *(a) Number of organoid cells sequenced from each layer of TRACER. (b) Number of genes detected in organoid cells from each layer of TRACER. (c) Percentage of reads from mitochondrial genes in organoid cells from each layer of TRACER. (d) Proportion of organoid cells classified in each phase of the cell cycle from each layer of TRACER. (e) Single cell expression of cell type marker genes in organoid cells from all layers of TRACER, with uniform identification of ductal 2 cells (KRT19) confirming the identity of organoid cells from all layers as malignant PDAC cells.*

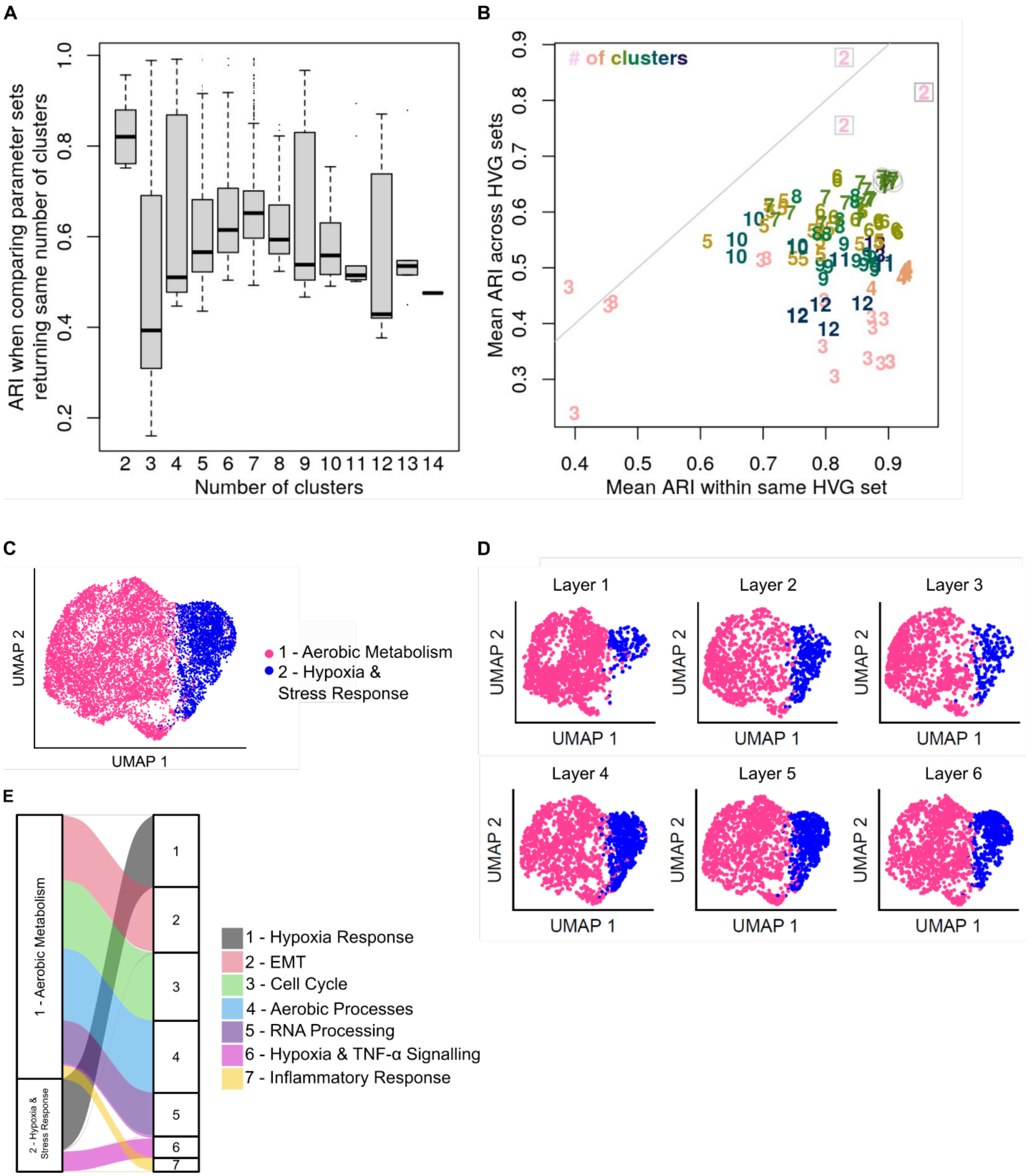


***Supplementary Figure S2: Optimization of cluster number shows organoid cells from TRACER can be grouped into 7 clusters.*** *(a) Adjusted Rand Index (ARI) for all pairs of cluster solutions with the same number of clusters. The 2 cluster solution is the most robust to changes in parameterization, but median ARI for 7 clusters is the next most robust. (b) Clustering was performed using different Highly Variable Gene (HVG) sets as input. Comparing ARIs within each set of HVGs versus across HVG inputs, the 2 cluster solutions (boxed) were most robust to changes in HVG input. The 7 cluster solution (circled) was the best-performing solution with more than two clusters. (c) UMAP reduction of the two cluster solution*

*with clusters annotated by GSEA enrichments (Cluster 1: "aerobic metabolism", Cluster 2: "hypoxia and stress response"). (d) UMAP reduction of the 2 cluster solution showing distribution of ODCs from Layer 1 (outer) to Layer 6 (inner) of TRACER2. The "hypoxia and stress response" cluster is better represented in the hypoxic inner layers of TRACER, consistent with known oxygen gradients in TRACER. (e) Mapping between 2 and 7 cluster solution highlighted by GSEA annotation. The variety of gene sets represented in the Cluster 1 highlighted the value of examining the 7 cluster solution.*

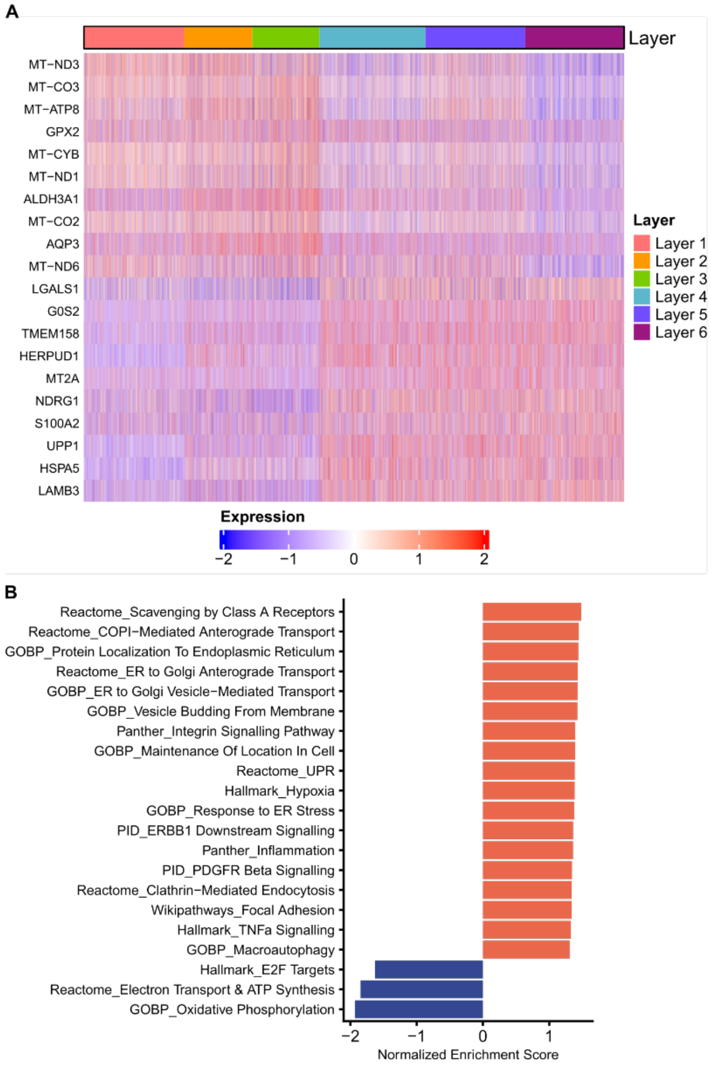


***Supplementary Figure S3: Gene sets related to the cellular response to hypoxia and stress show the most substantial enrichment across the TRACER layers.*** *(A) Heatmap of genes with the greatest positive or negative slope coefficient (top 10 each) for the linear regression of gene expression across the layers. Robust heterogeneity with clear trends in expression are present from Layer 1 (outer) to Layer 6 (inner). (B) Normalized enrichment score (NES) for gene sets significantly enriched (p < 0.05) based on the slope coefficient of the linear regression, where positive NES (red) indicates increasing enrichment from Layer 1 (outer) to Layer 6 (inner) and a negative NES (blue) indicates decreasing enrichment from Layer 1 to Layer 6. GSEA revealed expected positive enrichment of hypoxia and stress response related gene sets and negative enrichment of gene sets related to aerobic respiration and cell cycle.*

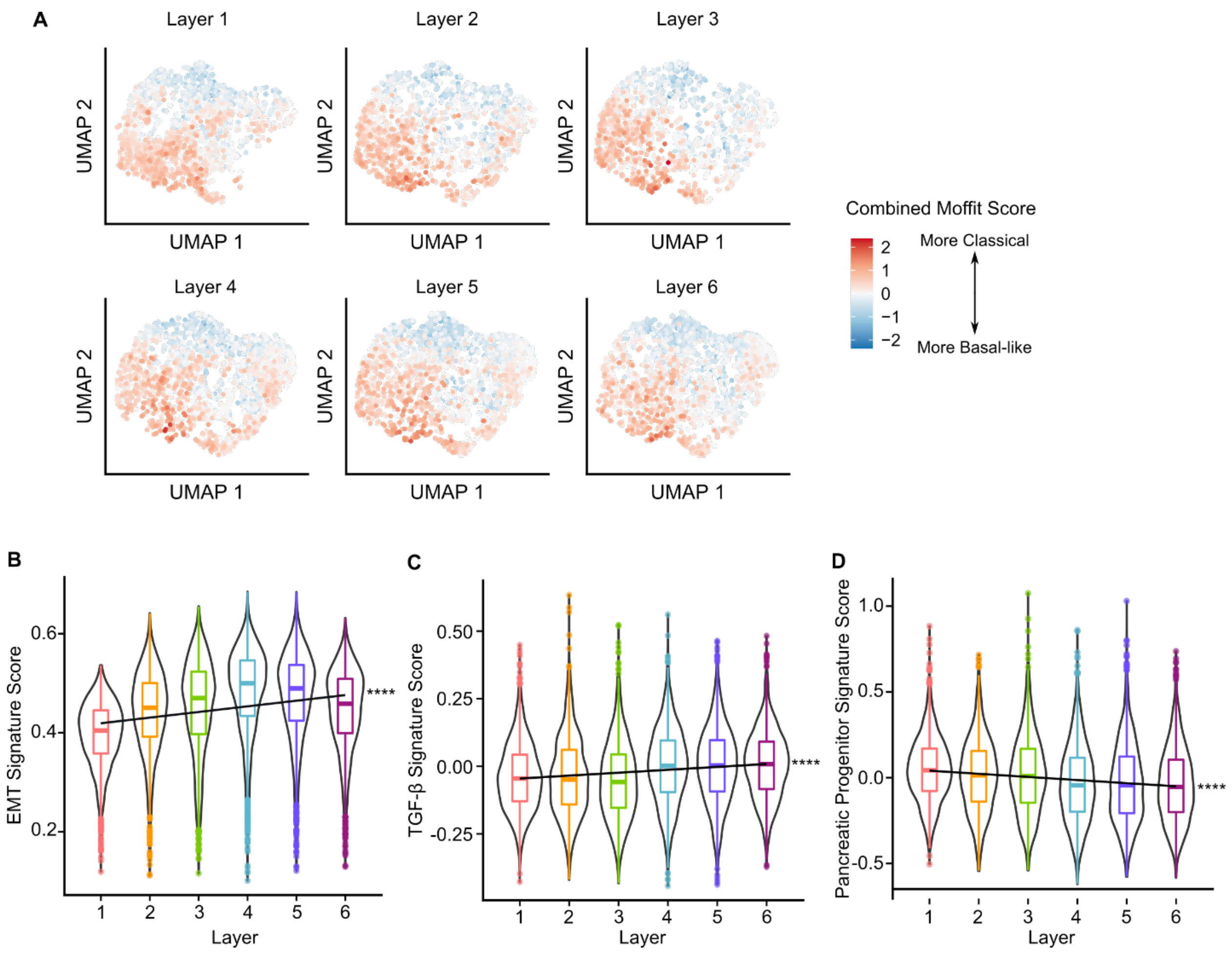


***Supplementary Figure S4: Microenvironmental gradients contribute to changes in the expression patterns of Moffit subtype signature genes and gene sets previously associated with the Moffit subtypes in organoid cells cultured in TRACER.*** *(a) UMAP reduction showing the combined Moffit score for organoid cells from each layer of TRACER. Changes in both the proportion of subtypes and the expression of subtype signature scores can be noted across the layers. (b)-(d) Gene signature expression scores for the (b) EMT (dbEMT consensus signature[1]), (c) TGF-β Receptor Signalling (Pathway Interaction Database), and (d) pancreatic progenitor signature[2] gene sets. The EMT and TGF-β receptor signalling scores exhibited a graded increase across the layers, consistent with their association with the basal-like subtype. Conversely, the pancreatic progenitor signature score exhibited a graded decreases across the layers, consistent with its association with the classical subtype (slope of the best fit line vs. 0, ****$p < 0.0001$).*

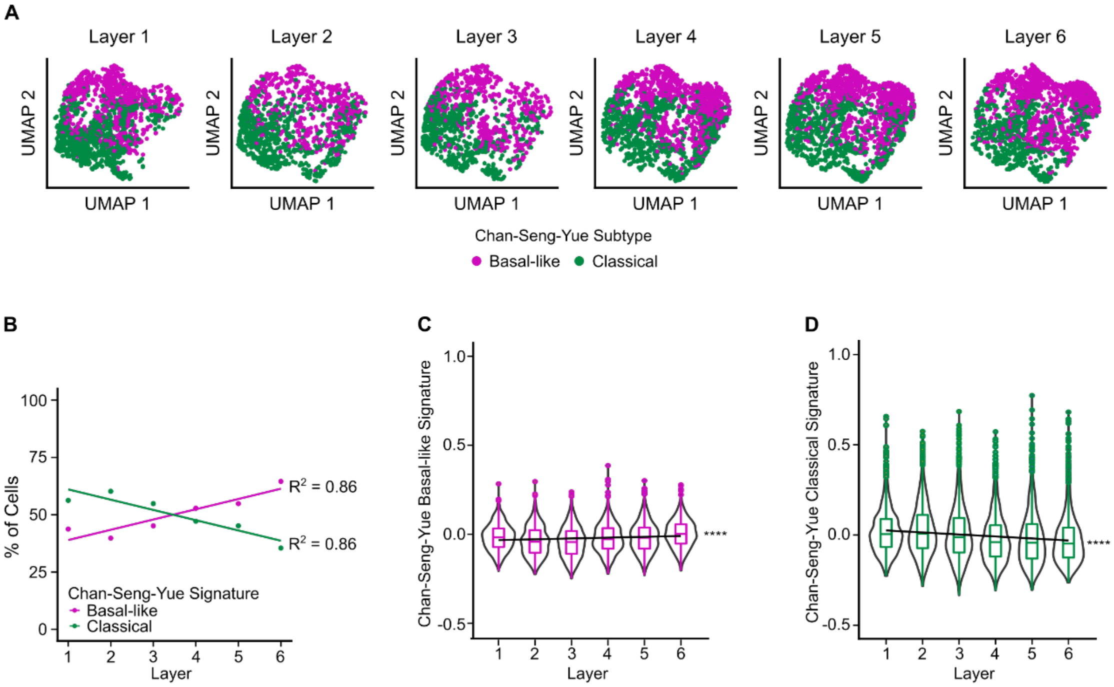


***Supplementary Figure S5: Microenvironmental gradients contribute to an increase in the proportion of basal-like organoid cells classified according to the Chan-Seng-Yue basal-like and classical gene signatures**. (A) UMAP plot showing the distribution of organoid cells classified by their maximal Chan-Seng-Yue signature scores in Layer 1 (outer) and Layer 6 (inner) of TRACER. (B) Quantification of organoid cell subtype classification proportions by layer shows a graded increase in the proportion of basal-like organoid cells towards the inner layers of TRACER. (C) The Chan-Seng-Yue basal-like signature score exhibits a significantly graded increase across TRACER layers (slope of the best fit line vs. 0, ****p< 0.0001) (D) The Chan-Seng-Yue classical signature score exhibits a significant graded decrease across TRACER layers (slope of the best fit line vs. 0, ****p<0.0001).*

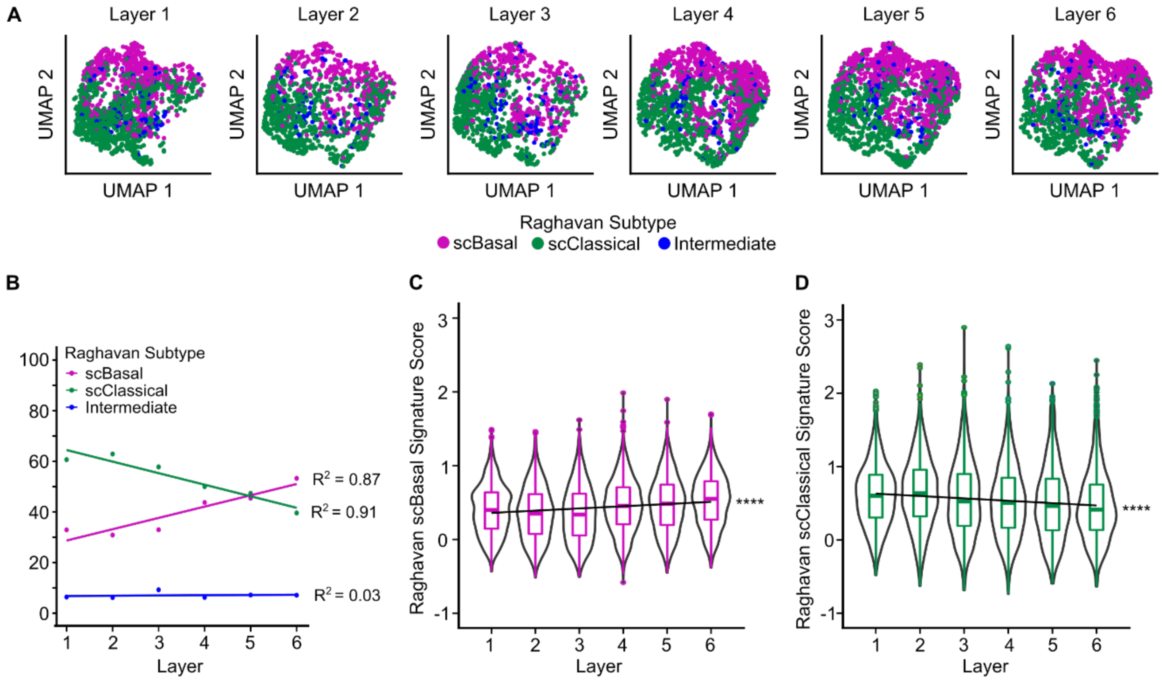


***Supplementary Figure S6: Microenvironmental gradients contribute to an increase in the proportion of basal-like organoid cells classified according to the Raghavan scBasal/scClassical gene signatures****.* *(A) UMAP plot showing the distribution of organoid cells classified by their maximal Raghavan signature scores in Layer 1 (outer) and Layer 6 (inner) of TRACER. (B) Quantification of organoid cell subtype classification proportions by layer shows a graded increase in the proportion of basal-like organoid cells towards the inner layers of TRACER. (C) The Raghavan scBasal signature score exhibits a significantly graded increase across TRACER layers (slope of the best fit line vs. 0, ****p< 0.0001) (D) The Raghavan scClassical signature score exhibits a significant graded decrease across TRACER layers (slope of the best fit line vs. 0, ****p<0.0001).*

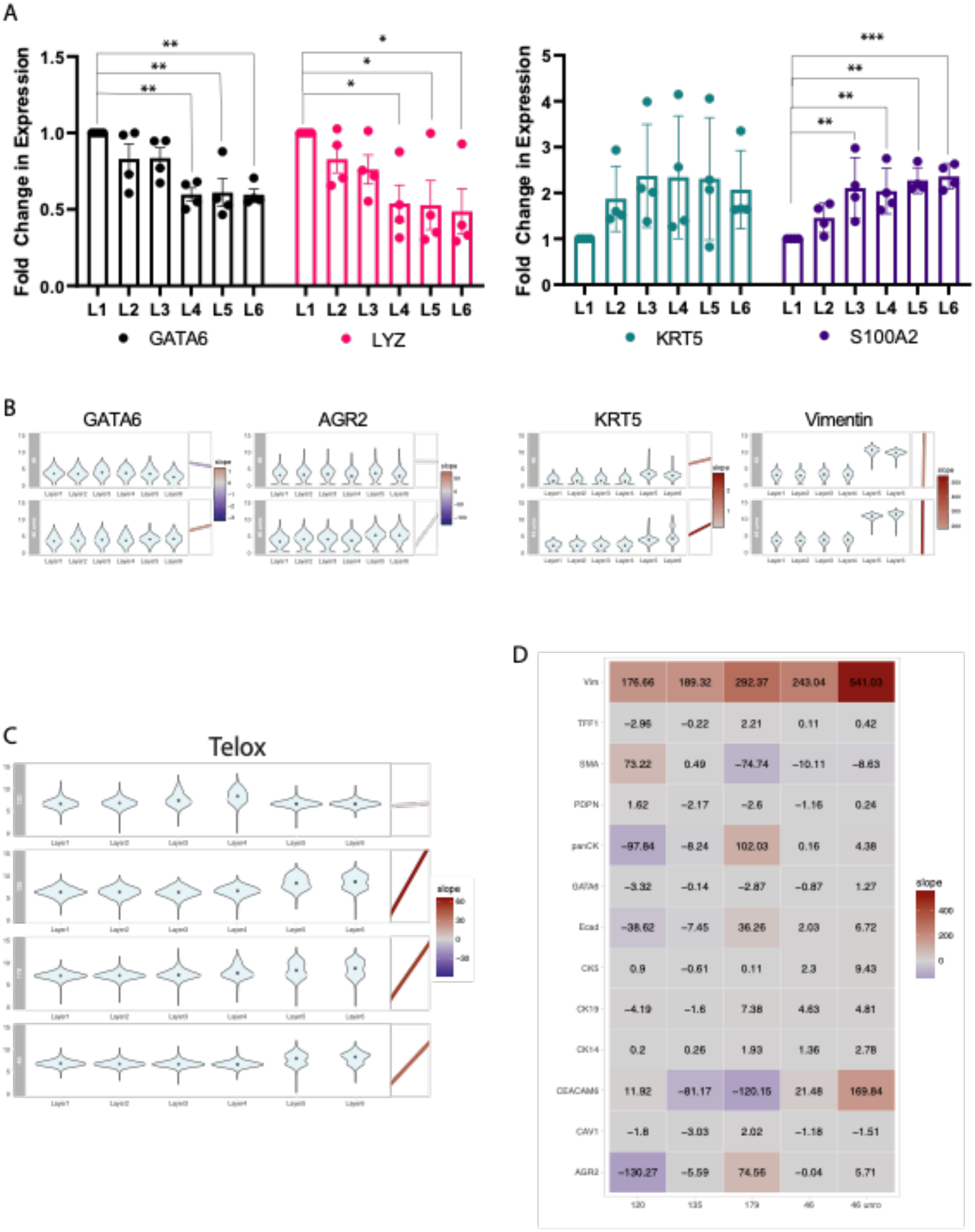

A
Fold Change in Expression
L1 L2 L3 L4 L5 L6
GATA6
LYZ
KRT5
S100A2
B
GATA6
AGR2
KRT5
Vimentin
C
Telox
D
Vim 176.66 189.32 292.37 243.04 541.03
TFF1 −2.96 −0.22 2.21 0.11 0.42
SMA 73.22 0.49 −74.74 −10.11 −8.63
PDPN 1.62 −2.17 −2.6 −1.16 0.24
panCK −97.84 −8.24 102.03 0.16 4.38
GATA6 −3.32 −0.14 −2.87 −0.87 1.27
Ecad −38.62 −7.45 36.26 2.03 6.72
CK5 0.9 −0.61 0.11 2.3 9.43
CK19 −4.19 −1.6 7.38 4.63 4.81
CK14 0.2 0.26 1.93 1.36 2.78
CEACAM6 11.92 −81.17 −120.15 21.48 169.84
CAV1 −1.8 −3.03 2.02 −1.18 −1.51
AGR2 −130.27 −5.59 74.56 −0.04 5.71
120 135 179 46 46 unro
slope

***Supplementary Figure S7: Validation of scRNA-seq in independent experiments with PPTO.46 organoid cells and exploration in new organoid lines.*** *(A) Gene expression of GATA6, LYZ (classical), KRT5 and S100A2 (basal-like) in PPTO.46 organoid cells in TRACER. Trends in classical and basal-like marker genes are consistent with scRNA-seq data, with decreases in classical marker gene expression and increases in S100A2 across the layers (4 independent experiments with n=3 TRACERs, one-way ANOVA with Dunnett's posthoc *p < 0.05, **p < 0.01, ***p < 0.001. ). (B)Assessment of the reversibility of the transcriptional change after reoxygenation using CyTOF. Measure of the expression of classical markers (GATA6 and AGR2) and basal markers (CK5 and Vimentin) in TRACER seeded with PPTO.46 without or with a 24h unrolled period after being rolled for 24h. Violin plots summarize the metal intensity of each cell and dark dots indicates the mean expression per layer for each organoid line and marker. The inserts on the right side of the violin plots indicates the best fit slope, lines are color coded by slope values (blue corresponding to negative slopes and red to positive slopes). (C)Measure of Telox signal as a surrogate for hypoxia using CyTOF. The level of TELOX in TRACER seeded with different organoid lines (organoid lines indicated in grey). Violin plots summarize the Telox intensity of each cell and dark dots indicates the mean expression per layer for each organoid line. The inserts on the right side of the violin plots indicates the best fit slope, lines are color coded by slope values (blue corresponding to negative slopes and red to positive slopes). (D) Summary for all markers of the measured slopes from layer 1 to layer 6 in the different TRACER seeded with the different organoid lines. Values indicate the measure slopes.*

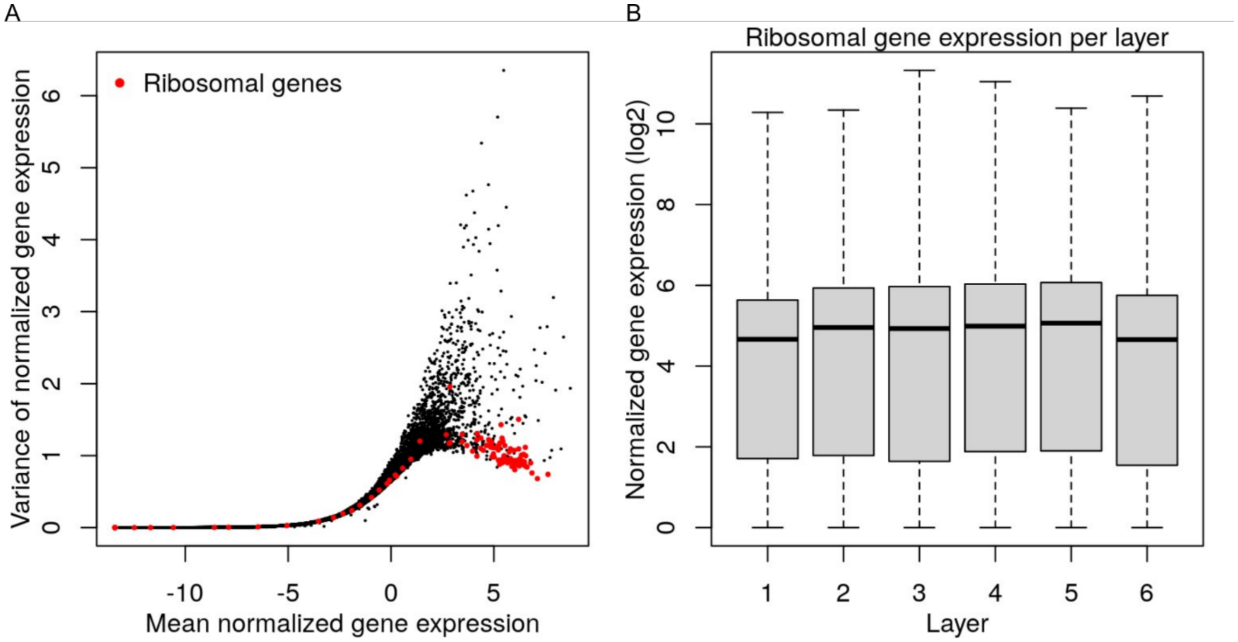


***Supplementary Figure S8: Appropriate normalization of gene expression from scRNA-seq dataset is verified by consistent ribosomal gene expression across the layers of TRACER.*** *(a) Scatterplot showing the mean-variance relationship of normalized gene expression for all genes across single-cell libraries from all layers. Ribosomal genes are highlighted in red. (b) Boxplots summarizing single-cell expression of ribosomal genes per layer.*

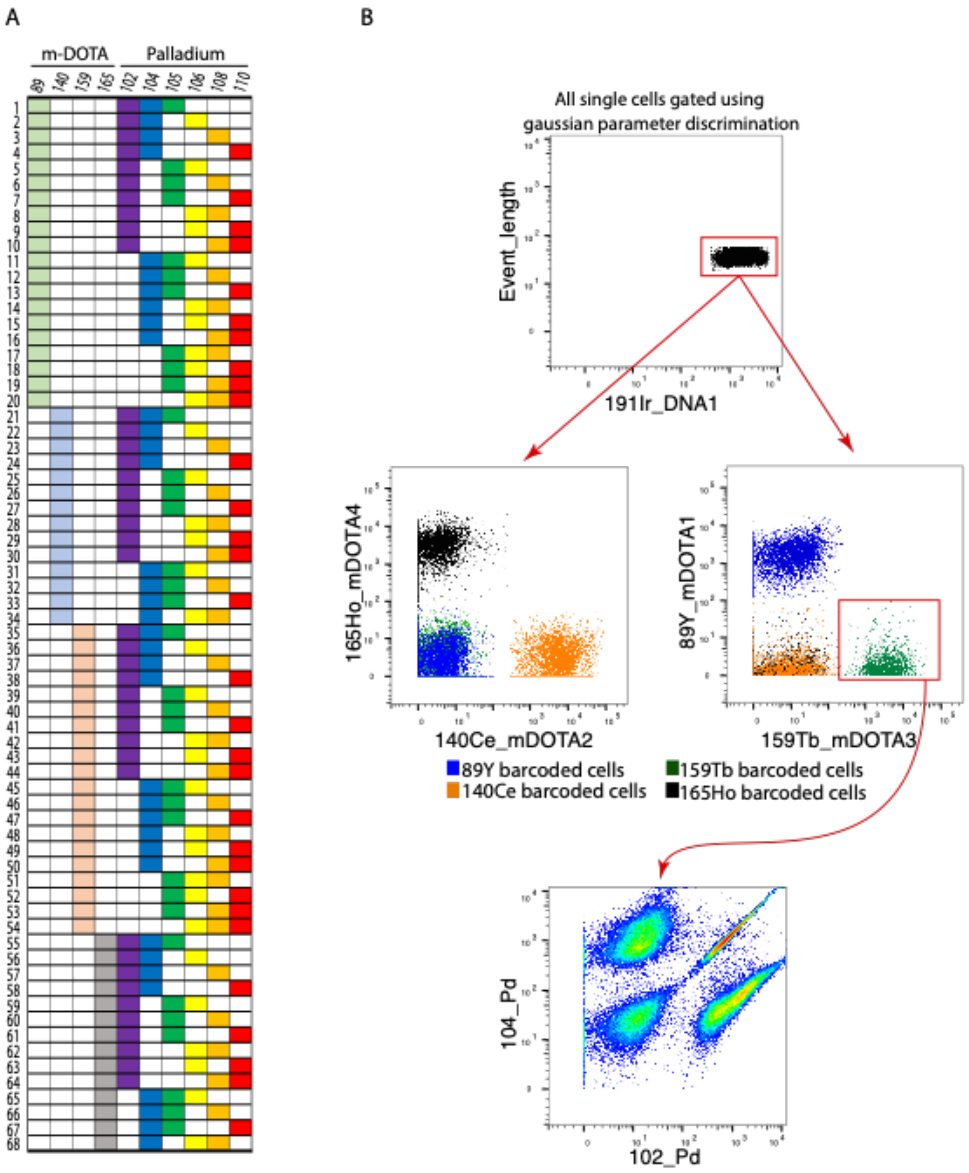


**Supplementary Figure S9: CyTOF barcoding and debarcoding strategy to allow for simultaneous measurement of 68 samples with a PDAC specific antibody panel.** (A) Barcoding map indicating the unique combination of metal use for the barcoding of individual samples. Barcoding was performed by combination of m-DOTA (4 column on the left isotopes: Y89, Ce140, Tb159 and Ho165) and palladium mix from the Cell-ID plex 20. (B) Gating strategy for debarcoding after acquisition. Using gaussian parameter discrimination, single cells were isolated. Using the four m-DOTA channels, cells were split in four .fcs files containing all cells barcoded with each of the m-DOTA. Each file was then processed using the debarcoder included in the CyTOF XT software. Debarcoding parameters were set to max manhaolis distance at 30 and minimum separation to 0.2.

**Supplementary Table S1: Details of the significantly enriched gene sets shown in Fig. 1E.**

| Heatmap Label | Gene Set Enriched on GSEA | Cluster | Adjusted p-value | Normalized Enrichment Score (NES) |
|---|---|---|---|---|
| Aerobic Respiration | AEROBIC RESPIRATION \| GOBP \| GO:0009060 | 1 | 7.16E-17 | -2.726069003 |
| Oxidative Phosphorylation | OXIDATIVE PHOSPHORYLATION \| GOBP \| GO:0006119 | 1 | 4.24E-15 | -2.782134219 |
| Aerobic ETC | AEROBIC ELECTRON TRANSPORT CHAIN \| GOBP \| GO:0019646 | 1 | 5.66E-12 | -2.648397377 |
| Response to ER Stress | RESPONSE TO ENDOPLASMIC RETICULUM STRESS \| GOBP \| GO:0034976 | 1 | 2.94E-06 | 1.576195345 |
| Unfolded Protein Response | UNFOLDED PROTEIN RESPONSE (UPR) \| REACTOME DATABASE ID RELEASE 77 \| 381119 | 1 | 1.57E-05 | 1.66811935 |
| Hypoxia | HALLMARK_HYPOXIA \| MSIGDB_C2 | 1 | 0.00080272 | 1.552155252 |
| E2F Targets | HALLMARK_E2F_TARGETS \| MSIGDB_C2 | 1 | 0.00138698 | -1.839336982 |
| Integrin Signalling | INTEGRIN SIGNALLING PATHWAY \| PANTHER PATHWAY \| P00034 | 1 | 0.00190522 | 1.531993854 |
| Cell Junction Organization | CELL JUNCTION ORGANIZATION \| REACTOME \| R-HSA-446728.2 | 1 | 0.00649501 | 1.596238356 |
| Aerobic ETC | RESPIRATORY ELECTRON TRANSPORT, ATP SYNTHESIS BY CHEMIOSMOTIC COUPLING, AND HEAT PRODUCTION BY UNCOUPLING PROTEINS. \| REACTOME \| R-HSA-163200.1 | 2 | 6.55E-21 | -2.754763197 |
| Aerobic Respiration | AEROBIC RESPIRATION \| GOBP \| GO:0009060 | 2 | 2.80E-16 | -2.62452836 |
| Oxidative Phosphorylation | HALLMARK_OXIDATIVE_PHOSPHORYLATION \| MSIGDB_C2 | 2 | 1.49E-15 | -2.467102466 |
| Response to ER Stress | RESPONSE TO ENDOPLASMIC RETICULUM STRESS \| GOBP \| GO:0034976 | 2 | 1.40E-07 | -2.003852799 |
| ECM Organization | EXTRACELLULAR MATRIX ORGANIZATION \| REACTOME DATABASE ID RELEASE 77 \| 1474244 | 2 | 1.61E-07 | 1.729657892 |
| mRNA Splicing | MRNA SPLICING \| REACTOME \| R-HSA-72172.3 | 2 | 2.11E-07 | -1.955965548 |
| Integrin Signalling | INTEGRIN SIGNALLING PATHWAY \| PANTHER PATHWAY \| P00034 | 2 | 2.90E-07 | 1.732019401 |
| G1S Transition | G1 S TRANSITION \| REACTOME DATABASE ID RELEASE 77 \| 69206 | 2 | 3.92E-07 | -2.171207998 |
| E2F Targets | HALLMARK_E2F_TARGETS \| MSIGDB_C2 | 2 | 8.40E-07 | -2.172974554 |
| Cell-cell Adhesion | CELL-CELL ADHESION \| GOBP \| GO:0098609 | 2 | 1.44E-06 | 1.662359128 |
| G2M Checkpoint | G2 M CHECKPOINTS \| REACTOME \| R-HSA-69481.3 | 2 | 1.47E-05 | -2.01523991 |
| Cell Cycle Checkpoints | CELL CYCLE CHECKPOINTS \| REACTOME \| R-HSA-69620.2 | 2 | 1.47E-05 | -1.77009682 |
| Cell Junction Organization | CELL JUNCTION ORGANIZATION \| GOBP \| GO:0034330 | 2 | 7.11E-05 | 1.56579086 |
| EMT | HALLMARK_EPITHELIAL_MESENCHYMAL_TRANSITION \| MSIGDB_C2 | 2 | 8.28E-05 | 1.704502194 |
| Unfolded Protein Response | HALLMARK_UNFOLDED_PROTEIN_RESPONSE \| MSIGDB_C2 | 2 | 0.00012793 | -1.9691394 |
| G2M Checkpoint | HALLMARK_G2M_CHECKPOINT \| MSIGDB_C2 | 2 | 0.00099563 | -1.800889492 |
| Chromosome Segregation | CHROMOSOME SEGREGATION \| GOBP \| GO:0007059 | 2 | 0.0020303 | -1.605974792 |
| Cell-matrix Adhesion | CELL-MATRIX ADHESION \| GOBP \| GO:0007160 | 2 | 0.00276917 | 1.631664242 |
| RNA Transcription | RNA POLYMERASE II TRANSCRIPTION TERMINATION \| REACTOME \| R-HSA-73856.4 | 2 | 0.00343939 | -1.866138723 |
| Antigen Presentation | ANTIGEN PRESENTATION: FOLDING, ASSEMBLY AND PEPTIDE LOADING OF CLASS I MHC \| REACTOME \| R-HSA-983170.3 | 2 | 0.007106 | -1.955770843 |
| DNA Replication | NUCLEAR DNA REPLICATION \| GOBP \| GO:0033260 | 2 | 0.00736344 | -1.916724196 |
| E2F Targets | HALLMARK_E2F_TARGETS \| MSIGDB_C2 | 3 | 1.94E-27 | 1.86621618 |

| | | | | |
|---|---|---|---|---|
| G2M Checkpoint | HALLMARK_G2M_CHECKPOINT \| MSIGDB_C2 | 3 | 2.33E-20 | 1.81286917 |
| Chromosome Segregation | CHROMOSOME SEGREGATION \| GOBP \| GO:0007059 | 3 | 2.33E-20 | 1.722762066 |
| Mitotic Prometaphase | MITOTIC PROMETAPHASE \| REACTOME \| R-HSA-68877.5 | 3 | 2.44E-15 | 1.654079291 |
| DNA Replication | DNA REPLICATION \| GOBP \| GO:0006260 | 3 | 2.04E-12 | 1.665514878 |
| Cell Cycle Checkpoints | REGULATION OF MITOTIC CELL CYCLE PHASE TRANSITION \| GOBP \| GO:1901990 | 3 | 1.03E-11 | 1.563241759 |
| S Phase | S PHASE \| REACTOME \| R-HSA-69242.2 | 3 | 2.57E-10 | 1.581186983 |
| G1S Transition | G1 S TRANSITION \| REACTOME DATABASE ID RELEASE 77 \| 69206 | 3 | 5.12E-09 | 1.587108896 |
| Aerobic ETC | RESPIRATORY ELECTRON TRANSPORT, ATP SYNTHESIS BY CHEMIOSMOTIC COUPLING, AND HEAT PRODUCTION BY UNCOUPLING PROTEINS. \| REACTOME \| R-HSA-163200.1 | 3 | 5.28E-08 | 1.569258938 |
| Oxidative Phosphorylation | HALLMARK_OXIDATIVE_PHOSPHORYLATION \| MSIGDB_C2 | 3 | 4.25E-07 | 1.490720537 |
| mRNA Splicing | MRNA SPLICING \| REACTOME \| R-HSA-72172.3 | 3 | 9.39E-07 | 1.452704878 |
| Aerobic Respiration | AEROBIC RESPIRATION \| GOBP \| GO:0009060 | 3 | 1.03E-06 | 1.544420452 |
| Unfolded Protein Response | PHOTODYNAMIC THERAPY-INDUCED UNFOLDED PROTEIN RESPONSE \| WIKIPATHWAYS_20210910 \| WP3613 \| HOMO SAPIENS | 3 | 2.94E-05 | -2.566806936 |
| EMT | HALLMARK_EPITHELIAL_MESENCHYMAL_TRANSITION \| MSIGDB_C2 | 3 | 0.00018754 | -2.0599697 |
| Cell Junction Organization | CELL JUNCTION ORGANIZATION \| REACTOME \| R-HSA-446728.2 | 3 | 0.00026532 | -2.107176666 |
| Aerobic ETC | RESPIRATORY ELECTRON TRANSPORT, ATP SYNTHESIS BY CHEMIOSMOTIC COUPLING, AND HEAT PRODUCTION BY UNCOUPLING PROTEINS. \| REACTOME \| R-HSA-163200.1 | 4 | 8.42E-14 | 2.342106124 |
| Aerobic Respiration | AEROBIC RESPIRATION \| GOBP \| GO:0009060 | 4 | 1.05E-13 | 2.334544039 |
| Oxidative Phosphorylation | OXIDATIVE PHOSPHORYLATION \| GOBP \| GO:0006119 | 4 | 1.07E-11 | 2.316880536 |
| EMT | HALLMARK_EPITHELIAL_MESENCHYMAL_TRANSITION \| MSIGDB_C2 | 4 | 6.08E-05 | -1.985583836 |
| Unfolded Protein Response | UNFOLDED PROTEIN RESPONSE (UPR) \| REACTOME DATABASE ID RELEASE 77 \| 381119 | 4 | 7.46E-05 | -1.94024823 |
| Integrin Signalling | INTEGRIN SIGNALLING PATHWAY \| PANTHER PATHWAY \| P00034 | 4 | 0.00020312 | -1.911261873 |
| Mitotic Prometaphase | MITOTIC PROMETAPHASE \| REACTOME \| R-HSA-68877.5 | 4 | 0.00020553 | -1.840805131 |
| Hypoxia | HALLMARK_HYPOXIA \| MSIGDB_C2 | 4 | 0.00107587 | -1.82237911 |
| Cell Junction Organization | CELL JUNCTION ORGANIZATION \| REACTOME \| R-HSA-446728.2 | 4 | 0.00430797 | -1.864700918 |
| Cell-matrix Adhesion | REGULATION OF CELL-SUBSTRATE ADHESION \| GOBP \| GO:0010810 | 4 | 0.00469684 | -1.807853069 |
| TNF-α Signalling | HALLMARK_TNFA_SIGNALING_VIA_NFKB \| MSIGDB_C2 | 4 | 0.00723195 | -1.722355315 |
| mRNA Processing | MRNA PROCESSING \| GOBP \| GO:0006397 | 5 | 0.00161319 | 2.168066345 |
| mRNA Translation | CYTOPLASMIC TRANSLATION \| GOBP \| GO:0002181 | 5 | 0.0020786 | 2.372474851 |
| TNF-α Signalling | HALLMARK_TNFA_SIGNALING_VIA_NFKB \| MSIGDB_C2 | 6 | 2.52E-08 | 1.582421009 |
| Hypoxia | HALLMARK_HYPOXIA \| MSIGDB_C2 | 6 | 0.0003637 | 1.515471214 |
| Oxidative Phosphorylation | OXIDATIVE PHOSPHORYLATION \| GOBP \| GO:0006119 | 6 | 0.00513162 | -1.967374971 |
| IFN-α Response | HALLMARK_INTERFERON_ALPHA_RESPONSE \| MSIGDB_C2 | 7 | 3.08E-05 | 1.882807987 |
| IFN-γ Response | HALLMARK_INTERFERON_GAMMA_RESPONSE \| MSIGDB_C2 | 7 | 1.19E-08 | 1.969170814 |
| Response to ER Stress | RESPONSE TO ENDOPLASMIC RETICULUM STRESS \| GOBP \| GO:0034976 | 7 | 8.83E-06 | 1.834679628 |
| Antigen Presentation | ANTIGEN PROCESSING AND PRESENTATION OF PEPTIDE ANTIGEN \| GOBP \| GO:0048002 | 7 | 7.40E-05 | 1.876868518 |

| | | | | |
|---|---|---|---|---|
| mRNA Splicing | MRNA SPLICING \| REACTOME \| R-HSA-72172.3 | 7 | 0.00475104 | -1.669155239 |
| DNA Replication | DNA REPLICATION \| GOBP \| GO:0006260 | 7 | 0.00683429 | -1.800839719 |
| E2F Targets | HALLMARK_E2F_TARGETS \| MSIGDB_C2 | 7 | 0.00822113 | -1.740503507 |

**Supplementary Table S2: Summary of statistical analysis for Figures 2-4 and S4-S7.** Bolded values denote statistical significance ($p < 0.05$).

<table>
<tr><th>Figure</th><th>n</th><th>Statistical Test</th><th>p value</th></tr>
<tr><td rowspan="2">2B – Cluster proportions</td><td rowspan="2">1 independent scRNA-seq experiment*</td><td>Chi-square test</td><td>p < 0.0001</td></tr>
<tr><td>Chi-square test for trend in proportions</td><td>Cluster 1 – Hypoxia Response, Cluster 3 -Cell Cycle, Cluster 4 – Aerobic Processes, Cluster 6 – Hypoxia/TNF-α: p < 0.0001<br>Cluster 2 – EMT: 0.3018<br>Cluster 7 – Inflammatory Response: 0.0011</td></tr>
<tr><td>2C – Cluster marker gene expression</td><td>1 independent scRNA-seq experiment*</td><td>Linear regression, slope of the best fit line vs. 0</td><td>DDIT3, ITGB1, MKI67, PDK4, SF3A2, TNFAIP3: p < 0.0001<br>IRF1: p = 0.3550</td></tr>
<tr><td>2D – Hypoxia Score</td><td>1 independent scRNA-seq experiment*</td><td>Linear regression, slope of the best fit line vs. 0</td><td>p < 0.0001</td></tr>
<tr><td>3C – Gemcitabine sensitivity score</td><td>1 independent scRNA-seq experiment*</td><td>Linear regression, slope of the best fit line vs. 0</td><td>p < 0.0001</td></tr>
<tr><td>3D – Basal-like signature score</td><td>1 independent scRNA-seq experiment*</td><td>Linear regression, slope of the best fit line vs. 0</td><td>p < 0.0001</td></tr>
<tr><td>3E – Classical signature score</td><td>1 independent scRNA-seq experiment*</td><td>Linear regression, slope of the best fit line vs. 0</td><td>p < 0.0001</td></tr>
<tr><td>3G – Hypoxia response pathway signature correlation</td><td>1 independent scRNA-seq experiment*</td><td>Spearman correlation coefficient</td><td>Signature comparisons<br>Basal-like/HIF-1α, basal-like/mTOR, classical/mTOR: p = 0<br>Classical/HIF-1α: p = 1.58 x $10^{-5}$<br>Classical/UPR: p = 1.93x$10^{-95}$<br>Basal-like/UPR: p = 0.0094</td></tr>
<tr><td>4A – Hypoxia chamber comparison, gene expression</td><td>Layer 1 and Layer 6: 4 independent experiments, each an average of 3 technical replicates; 0.2% pO2: 5 independent experiments, each an</td><td>One-sample t-test of actual mean vs. theoretical mean of 1 (normoxic</td><td><table>
<tr><th></th><th>L1</th><th>L6</th><th>0.2% pO2</th></tr>
<tr><td>GATA6</td><td>0.2102</td><td>0.0222</td><td>0.0173</td></tr>
<tr><td>LYZ</td><td>0.6353</td><td>0.0370</td><td><0.0001</td></tr>
<tr><td>KRT5</td><td>0.3930</td><td>0.1539</td><td>0.0382</td></tr>
<tr><td>S100A2</td><td>0.1854</td><td>0.0484</td><td>0.0803</td></tr>
</table></td></tr>
</table>

<table>
<tr><td></td><td>average of 3 technical replicates</td><td>control as reference)</td><td></td></tr>
<tr><td>4B – PPTO46 GATA6 CyTOF</td><td>4 independent experiments, each an average of 2 technical replicates for both TRACER2 and single layer controls</td><td>TRACER2: Linear regression, slope of the trend line vs. 0; Normoxia vs 0.2% O2: t-test</td><td>TRACER2: p = 0.0257<br>Normoxia vs 0.2% $pO_2$: p = 0.0066</td></tr>
<tr><td rowspan="2">4C – PPTO120 qPCR at 24h</td><td rowspan="2">3 independent experiments, each an average of 3 technical replicates</td><td>One-way ANOVA with Dunnett’s posthoc multiple comparison test (compared to L1)</td><td><table>
<tr><td>L1 vs</td><td>L2</td><td>L3</td><td>L4</td><td>L5</td><td>L6</td></tr>
<tr><td>GATA6</td><td>0.0766</td><td>0.0008</td><td>0.0003</td><td>0.0001</td><td><0.0001</td></tr>
<tr><td>LYZ</td><td>0.9466</td><td>0.1959</td><td>0.2476</td><td>0.1052</td><td>0.0132</td></tr>
<tr><td>KRT5</td><td>0.9846</td><td>0.5808</td><td>0.6220</td><td>0.8504</td><td>0.6560</td></tr>
<tr><td>S100A2</td><td>0.9049</td><td>0.8900</td><td>0.3306</td><td>0.6294</td><td>0.7921</td></tr>
</table></td></tr>
<tr><td>Linear regression, slope of the best fit line vs. 0</td><td>GATA6: p < 0.0001<br>LYZ: p = 0.0005<br>KRT5: p = 0.2085<br>S100A2: p = 0.2077</td></tr>
<tr><td>4D – PPTO120 GATA6 CyTOF</td><td>4 independent experiments, each an average of n=2-3 technical replicates for both TRACER2 and single-layer controls; 1 TRACER2 sample removed as a significant outlier</td><td>TRACER2: Linear regression, slope of the trend line vs. 0; Normoxia vs 0.2% O2: t-test</td><td>TRACER2: p = 0.0092<br>Normoxia vs 0.2% $pO_2$: p = 0.0145</td></tr>
<tr><td>4E – PPTO46 qPCR at 24h Rolled and 24h Rolled/24h Unrolled</td><td>24h rolled: 4 independent experiments, each an average of 3 technical replicates; 24h rolled/24h unrolled: 3 independent experiments each an average of 3 technical replicates</td><td>Linear regression, slope of the trend line vs. 0</td><td><table>
<tr><td></td><td>24h Rolled</td><td>24h Rolled/24h Unrolled</td></tr>
<tr><td>GATA6</td><td>< 0.0001</td><td>0.7861</td></tr>
<tr><td>LYZ</td><td>0.0005</td><td>0.0292</td></tr>
<tr><td>KRT5</td><td>0.1198</td><td>0.0607</td></tr>
<tr><td>S100A2</td><td>< 0.0001</td><td>0.6912</td></tr>
</table></td></tr>
<tr><td>S4B – EMT</td><td>1 independent scRNA-seq experiment*</td><td>Linear regression, slope of the</td><td>p < 0.0001</td></tr>
</table>

<table>
<tr><td>Signature Score</td><td></td><td>best fit line vs. 0</td><td></td></tr>
<tr><td>S4C – TGF-β Signature Score</td><td>1 independent scRNA-seq experiment*</td><td>Linear regression, slope of the best fit line vs. 0</td><td>p < 0.0001</td></tr>
<tr><td>S4D – Pancreatic Progenitor Signature Score</td><td>1 independent scRNA-seq experiment*</td><td>Linear regression, slope of the best fit line vs. 0</td><td>p < 0.0001</td></tr>
<tr><td>S5C – Chan-Seng-Yue Basal-like signature score</td><td>1 independent scRNA-seq experiment*</td><td>Linear regression, slope of the best fit line vs. 0</td><td>p < 0.0001</td></tr>
<tr><td>S5D – Chan-Seng-Yue Classical signature score</td><td>1 independent scRNA-seq experiment*</td><td>Linear regression, slope of the best fit line vs. 0</td><td>p < 0.0001</td></tr>
<tr><td>S6C – Raghavan scBasal signature score</td><td>1 independent scRNA-seq experiment*</td><td>Linear regression, slope of the best fit line vs. 0</td><td>p < 0.0001</td></tr>
<tr><td>S6D – Raghavan scClassical signature score</td><td>1 independent scRNA-seq experiment*</td><td>Linear regression, slope of the best fit line vs. 0</td><td>p < 0.0001</td></tr>
<tr><td>S7A – PPTO46 qPCR at 24h</td><td>4 independent experiments, each an average of 3 technical replicates</td><td>One-way ANOVA with Dunnett’s posthoc multiple comparison test (compared to L1)</td><td><table>
<tr><td>L1 vs</td><td>L2</td><td>L3</td><td>L4</td><td>L5</td><td>L6</td></tr>
<tr><td>GATA6</td><td>0.2742</td><td>0.2969</td><td>0.0018</td><td>0.0023</td><td>0.0017</td></tr>
<tr><td>LYZ</td><td>0.7349</td><td>0.4607</td><td>0.0426</td><td>0.0372</td><td>0.0220</td></tr>
<tr><td>KRT5</td><td>0.6242</td><td>0.2349</td><td>0.2497</td><td>0.2693</td><td>0.4434</td></tr>
<tr><td>S100A2</td><td>0.3804</td><td>0.0042</td><td>0.0070</td><td>0.0012</td><td>0.0006</td></tr>
</table></td></tr>
<tr><td>S7B – PPTO120 Telox CyTOF</td><td>4 independent experiments, each an average of n=2-3 technical replicates for both TRACER2 and single-layer</td><td>TRACER2: Linear regression, slope of the best fit line vs. 0;</td><td>TRACER2: p = 0.0003<br>Normoxia vs 0.2% $pO_2$: p < 0.0001</td></tr>
</table>

<table>
<tr><td></td><td>controls; 1 TRACER2 sample removed as a significant outlier</td><td>Normoxia vs 0.2% O2: t-test</td><td colspan="6"></td></tr>
<tr><td>S7C – PPTO120 IdU CyTOF</td><td>4 independent experiments, each an average of n=2-3 technical replicates for both TRACER2 and single-layer controls; 1 TRACER2 sample removed as a significant outlier</td><td>TRACER2: One-way ANOVA with Dunnett’s posthoc multiple comparison test (compared to L1)</td><td colspan="6">L1 vs L2: 0.4058<br>L1 vs L3: 0.0961<br>L1 vs L4: 0.0615<br>L1 vs L5: 0.0322<br>L1 vs L6: 0.0283</td></tr>
<tr><td rowspan="6">S7D – PPTO120 qPCR at 24h</td><td rowspan="6">3 independent experiments, each an average of 3 technical replicates</td><td rowspan="5">One-way ANOVA with Dunnett’s posthoc multiple comparison test (compared to L1)</td><td>L1 vs</td><td>L2</td><td>L3</td><td>L4</td><td>L5</td><td>L6</td></tr>
<tr><td>CHOP</td><td>0.9998</td><td>>0.999</td><td>0.7272</td><td>0.0203</td><td>0.0049</td></tr>
<tr><td>CA9</td><td>0.5884</td><td>0.9989</td><td>>0.999</td><td>0.8866</td><td>0.4595</td></tr>
<tr><td>REDD1</td><td>0.9549</td><td>0.9099</td><td>0.1380</td><td>0.0027</td><td>0.0022</td></tr>
<tr><td>ERDJ4</td><td>>0.999</td><td>>0.999</td><td>0.9986</td><td>0.7550</td><td>0.8878</td></tr>
<tr><td>Linear regression, slope of the best fit line vs. 0</td><td colspan="6">CHOP: p < 0.0001<br>CA9: p = 0.0334<br>REDD1: p < 0.0001<br>ERDJ4: p = 0.1862</td></tr>
</table>

*scRNA-seq experiment cell count by layer: Layer 1 = 2041 cells, Layer 2 = 1385 cells, Layer 3 = 1362 cells, Layer 4 = 2154 cells, Layer 5 = 2026 cells, Layer 6 = 2010 cells

**Supplementary Table S3: Details of the gene sets shown in Fig. 2F that show significantly graded enrichment in the microenvironment in TRACER.** A positive normalized enrichment score (NES) indicates that the gene set is increasingly enriched towards Layer 6 (inner). The truncated leading edge highlights the genes that most strongly contributed to the gene set being enriched.

| | Adjusted p-value | Normalized Enrichment Score (NES) | Truncated Leading Edge |
|---|---|---|---|
| HALLMARK_E2F_TARGETS \| MSIGDB_C2 | 5.67E-21 | -2.91054 | NOP56, DCTPP1, PCNA, UBE2T, MKI67, PA2G4 |
| ER TO GOLGI ANTEROGRADE TRANSPORT \| REACTOME \| R-HSA-199977.4 | 7.89E-21 | 2.136244 | KDELR1, TMED9, ARF4, ARF1, KDELR2, COPE |
| RESPONSE TO ENDOPLASMIC RETICULUM STRESS \| GOBP \| GO:0034976 | 4.30E-18 | 1.971812 | P4HB, CALR, HSPA5, HM13, SERP1, PDIA3 |
| **RESPIRATORY ELECTRON TRANSPORT**, ATP SYNTHESIS BY CHEMIOSMOTIC COUPLING, AND HEAT PRODUCTION BY UNCOUPLING PROTEINS. \| REACTOME \| R-HSA-163200.1 | 3.57E-15 | -2.68132 | MT-ND3, MT-CO3, MT-CYB, MT-ATP8, MT-CO2, MT-ND1 |
| OXIDATIVE PHOSPHORYLATION \| GOBP \| GO:0006119 | 1.82E-13 | -2.79251 | MT-ND3, MT-CO3, MT-CYB, MT-ATP8, MT-CO2, MT-ND1 |
| AEROBIC RESPIRATION \| GOBP \| GO:0009060 | 1.82E-13 | -2.66574 | MT-ND3, MT-CO3, MT-CYB, MT-ATP8, MT-CO2, MT-ND1 |
| HALLMARK_HYPOXIA \| MSIGDB_C2 \| HALLMARK_HYPOXIA | 2.16E-13 | 1.966614 | BNIP3L, BTG1, ANGPTL4, PRDX5, PPP1R15A, DDIT4 |
| ERBB1 DOWNSTREAM SIGNALING \| PATHWAY INTERACTION DATABASE NCI-NATURE CURATED DATA | 8.21E-11 | 1.962483 | RAC1, BRK1, ARPC1B, ARF4, CAPN2, YWHAZ |
| HALLMARK_MYC_TARGETS_V2 \| MSIGDB_C2 | 2.47E-10 | -2.77053 | NOP56, DCTPP1, PA2G4, NOLC1, SORD, AIMP2 |
| REGULATION OF ACTIN FILAMENT POLYMERIZATION \| GOBP \| GO:0030833 | 8.58E-10 | 1.899435 | BRK1, ARPC1B, ARF1, MTPN, ARPC1A, ARPC5 |
| VESICLE BUDDING FROM MEMBRANE \| GOBP \| GO:0006900 | 1.61E-09 | 2.062814 | TMED9, ANXA2, VAPA, TMED2, SEC24D, TMED10 |
| PROTEIN LOCALIZATION TO ENDOPLASMIC RETICULUM \| GOBP \| GO:0070972 | 1.81E-09 | 2.06479 | KDELR1, HSPA5, PPP1R15A, SEC61G, KDELR2, SEC61B |
| ACTIVATION OF THE PRE-REPLICATIVE COMPLEX \| REACTOME \| R-HSA-68962.4 | 1.98E-08 | -2.68345 | MCM3, GMNN, MCM6, MCM7, MCM4, MCM2 |
| SIGNALING BY MET \| REACTOME \| R-HSA-6806834.2 | 2.43E-07 | 1.932155 | SPINT2, RAC1, LAMB3, SH3GL1, ITGB1, LAMC2 |
| AUTOPHAGOSOME ASSEMBLY \| GOBP \| GO:0000045 | 5.59E-07 | 1.937216 | MAP1LC3B, RAB7A, GABARAPL2, WIPI1, RAB1A, GABARAP |
| RAS PATHWAY \| PANTHER PATHWAY \| P04393 | 3.18E-07 | 1.916664 | RAC1, RHOA, ETS1, RHOC, CDC42, MAP2K1 |
| PHOTODYNAMIC THERAPY-INDUCED **UNFOLDED PROTEIN RESPONSE** \| WIKIPATHWAYS_20210910 \| WP3613 \| HOMO SAPIENS | 3.48E-06 | 1.966828 | PPP1R15A, DNAJC3, DDIT3, WARS, TRIB3, DNAJB9 |
| HALLMARK_EPITHELIAL_MESENCHYMAL_TRANSITION \| MSIGDB_C2 | 1.00E-05 | 1.690651 | PPIB, CALU, GADD45A, VEGFA, TIMP1, ITGB1 |
| SENESCENCE AND AUTOPHAGY IN CANCER \| WIKIPATHWAYS_20210910 \| WP615 \| HOMO SAPIENS | 1.47E-05 | 1.795689 | CEBPB, HMGA1, CDKN1A, GABARAPL2, GABARAP, MAP2K1 |
| PROGRAMMED CELL DEATH \| REACTOME \| R-HSA-5357801.2 | 1.61E-05 | 1.596605 | YWHAZ, CDH1, STK24, UBC, SDCBP, PSMD8 |
| EPHB-MEDIATED FORWARD SIGNALING \| REACTOME DATABASE ID RELEASE 77 \| 3928662 | 3.26E-05 | 1.917461 | RAC1, ARPC1B, RHOA, ARPC1A, CDC42, ARPC5 |
| PID_MTOR_4PATHWAY \| MSIGDB_C2 | 4.03E-05 | 1.822024 | RAC1, DDIT4, YWHAZ, PXN, RHEB, EIF4EBP1 |
| DEGRADATION OF THE EXTRACELLULAR MATRIX \| REACTOME \| R-HSA-1474228.4 | 0.000527 | 1.633608 | LAMB3, CAPN2, CAST, CDH1, TIMP1, LAMC2 |
| HIF-1-ALPHA TRANSCRIPTION FACTOR NETWORK \| PATHWAY INTERACTION DATABASE NCI-NATURE CURATED DATA | 0.000982 | 1.691797 | SLC2A1, NDRG1, ETS1, PGK1, HK2, LDHA |

| | | | |
|---|---|---|---|
| FATTY ACID BETA-OXIDATION \| GOBP \| GO:0006635 | 0.009994 | -1.70292 | HADH, MTLN, ACADM, MCAT, IVD, SLC27A2 |
| CELLULAR RESPONSE TO XENOBIOTIC STIMULUS \| GOBP \| GO:0071466 | 0.00128 | -1.71699 | ALDH3A1, CYP1B1, UGT1A6, GSTP1, CYB5B, UGT1A1 |
| TRYPTOPHAN METABOLISM \| WIKIPATHWAYS_20210910 \| WP465 \| HOMO SAPIENS | 0.006438 | -1.88694 | CYP1B1, DHCR24, CYP1A1, AOC1, ALDH3A2, DDC |
| STRESS GRANULE ASSEMBLY \| GOBP \| GO:0034063 | 0.006156 | 1.675285 | CIRBP, DYNC1H1, PRRC2C, BICD1, ATXN2L, CSDE1 |

**Supplementary Table S4: qPCR primers**

| Gene | Forward Primer (5' to 3') | Reverse Primer (5' to 3') |
|---|---|---|
| CA9 | CATCCTAGCCCTGGTTTTTGG | GCTCACACCCCCTTTGGTT |
| CHOP | GGAGCATCAGTCCCCCACTT | TGTGGGATTGAGGGTCACATC |
| ERdj4 | AAAATAAGAGCCCGGATGCT | CGCTTCTTGGATCCAGTGTT |
| GATA6 | GTGCCCAGACCACTTGCTAT | TGGAATTATTGCTATTACCAGAGC |
| KRT5 | GCTGCCTACATGAACAAGGTGG | ATGGAGAGGACCACTGAGGTGT |
| LYZ | ACTACAATGCTGGAGACAGAAGC | GCACAAGCTACAGCATCAGCGA |
| REDD1 | GGTCACTGAGCAGCTCGAA | CCTGGACAGCAGCAACAGT |
| RPL13a | CCGGGTTGGCTGGAAGTACC | CTTCTCGGCCTGTTTCCGTAG |
| S100A2 | TGCCAAGAGGGCGACAAGTTCA | AAGTCCACCTGCTGGTCACTGT |

**Supplementary Table S5: CyTOF antibodies**

| Target | Reference | Clone | Metal Conjug |
|---|---|---|---|
| Ecad | BD Biosciences 610182 | 36/E-Cadherin | In113 |
| Vim | Abcam ab193555 | EPR3776 | Nd142 |
| CEACAM6 | Abcam ab275033 | EPR23956-80 | Nd143 |
| CK5 | Abcam ab214586 | EP1601Y | Nd144 |
| CK14 | Abcam ab212547 | LL002 | Nd145 |
| CK19 | Sigma MABT913 | Troma-III | Nd146 |
| AGR2 | Cell Signaling 1551 | D9V2F | Sm149 |
| TFF1 | Cell Signaling 15571 | D2Y1J | Nd150 |
| GATA6 | Novus Biologicals AF1700 | AF1700 | Eu151 |
| CAV1 | Cell signaling 3267 | D46G3 | Tm169 |
| PDPN | Biolegend 337002 | NC-08 | Lu175 |
| panCK | Thermo Fisher MA1-12594 | C11 | Pt198 |
| SMA | Thermo Fisher 14-9760-82 | 1A4 | Bi209 |
